# Cholinergic recruitment of Chandelier cells during arousal

**DOI:** 10.1101/2025.01.18.633708

**Authors:** Emilio Martinez-Marquez, Santiago Reyes-Leon, Flores Garcia-San-Jose, Santiago Lopez-Begines, Jose L Nieto-Gonzalez, Pablo Garcia-Junco-Clemente

## Abstract

Chandelier cells (ChCs) are a highly specialized subtype of GABAergic interneurons and one of the most distinctive elements of cortical circuitry, exerting powerful and strategic control over pyramidal neuron output by selectively innervating their axon initial segment. They are particularly abundant in the prefrontal cortex, where cholinergic inputs modulate cognitive functions and shape ChC axonal development, but the way in which the cholinergic system—a master modulator of attention and arousal—regulates these cells in the adult brain has long remained unexplored. In this study, by employing an intersectional genetic strategy in adult mice, we reveal that ChCs in the secondary motor cortex are direct targets of cholinergic modulation. Through patch-clamp recordings and functional imaging, we demonstrate that acetylcholine persistently activates ChCs via heteromeric nicotinic receptors containing the β2 subunit, triggering robust depolarization and a significant increase in intrinsic excitability. This regulation does not rely on fast synaptic transmission; instead, it arises from a diffuse and sustained cholinergic signaling mode, orchestrated from the basal forebrain. Intriguingly, our in vivo observations show that ChC activity is positively correlated with behavioral markers of high arousal, such as locomotion and pupil dilation—a signature of activity that diminished upon the blockade of nicotinic receptors. Our findings strongly suggest that ChCs serve as a link between global arousal and local cortical control, thereby offering deeper insights into the mechanisms of state-dependent information processing.

## Introduction

The mammalian cerebral cortex is a highly complex structure consisting of excitatory and inhibitory neurons that form intricate networks essential for brain function. Excitatory pyramidal neurons (PNs), which constitute approximately 85–90% of all cortical neurons, are primarily responsible for transmitting information between cortical areas and relaying signals from the cortex to other brain regions (*1*, *2*). In contrast, inhibitory GABAergic interneurons, although comprising only 10-15% of cortical neurons, play a crucial role in the control of synaptic hyperexcitability and the fine-tuning synaptic integration, spike timing and temporal coordination within cortical circuits (*3–7*). These interneurons are traditionally classified based on morphological and electrophysiological properties, with key molecular markers such as parvalbumin, somatostatin, and the ionotropic serotonin receptor 5HT3a defining major subtypes (*2*, *7*, *8*). Despite advances in interneuron classification, significant heterogeneity exists within these groups, leaving the precise contributions of many interneuron subtypes unresolved.

Among GABAergic interneurons, chandelier cells (ChCs), also known as axo-axonic interneurons, are particularly notable due to their highly specialized connectivity (*9*). ChCs are a subclass of parvalbumin-expressing interneurons, although it has been reported that some ChCs may exhibit weak or negligible parvalbumin expression (*10*), underscoring the complexity of interneuron classification. These interneurons specifically target the axon initial segment (AIS) of PNs, a critical site for action potential initiation, allowing ChCs to exert significant influence on neuronal output and network activity (*11–14*).

Despite significant progress in targeting ChCs through recent genetic strategies, their precise functional contribution to cortical processing remains a subject of intense study. The availability of genetically encoded tools has shifted the focus toward a systematic manipulation of these interneurons in vivo (*10*, *13*, *15–21*). These tools have helped to demonstrate that ChCs integrate inputs from both local excitatory and inhibitory neurons as well as higher-order cortical regions, emphasizing their role as integrative hubs in cortical networks (*13*, *22*).

Acetylcholine (ACh) is a key neuromodulator in the central nervous system, critically involved in regulating arousal, attention, and learning by modulating neuronal excitability, synaptic integration, and circuit plasticity (*23–27*). The basal forebrain (BF) serves as the principal source of cholinergic input to the cortex, with projections primarily targeting superficial layers (*25*, *28*, *29*). Notably, ChCs reside at the layer 1–layer 2/3 (L1-L2/3) boundary, extending their dendritic branches within the lamina and sending a prominent dendrite upward into L1 (*10*, *30*). Indeed, prefrontal L1 contains one of the highest laminar densities of cholinergic axons and varicosities (*31*), suggesting that ChCs may be strongly influenced by the cholinergic system, which is critically involved in the regulation of brain network states. Supporting this notion, ChCs have been shown to receive direct synaptic input from the diagonal band (*13*). Elevated cholinergic tone characterizes high-arousal states, often reflected in behavioral readouts such as locomotion or pupil dilation (*32–35*). Specifically, ChCs exhibit heightened activity during high-arousal conditions, such as locomotion, across cortical regions including the hippocampus, visual cortex, and secondary motor area (M2) (*17*, *18*, *20*, *36*). Moreover, cholinergic signaling governs the axonal arborization of ChCs during development (*37*).

In this study, we explore whether ChCs may be positioned to act as hubs aligning local circuit activity with global cholinergic signals during specific cortical states. To address this, we employed an intersectional genetic approach (*38*) to selectively label these interneurons for comprehensive morphological, electrophysiological, and immunohistochemical characterization. By combining in vitro recordings and in vivo imaging with optogenetic manipulations, we sought to define the mechanisms of cholinergic modulation via nicotinic acetylcholine receptors (nAChRs). Our data demonstrate that M2 ChCs express functional heteromeric receptors and exhibit robust depolarization under cholinergic drive. Furthermore, ChC activity patterns are tightly coupled with key indicators of arousal, such as locomotion and pupil dilation, an association that is disrupted by in vivo nicotinic blockade. Given the proposed role of ChCs in coordinating cortical network activity, these findings raise the possibility that cholinergic modulation of ChC excitability could influence cortical dynamics during active behavioral states.

## Results

### Assessment of the *VipR2-Pvalb-Ai65* Mouse Model for Precise Targeting of Chandelier Cells

To investigate cholinergic modulation in ChCs, we first established a robust genetic targeting framework for their visualization and characterization. Following the intersectional strategy described by Tasic et al. (2018), we utilized the dual expression of *Vipr2* and *Pvalb* promoters to achieve high-precision labeling of this interneuron population. Specifically, CRE recombinase was driven by the *Vipr2* promoter, which is selectively expressed in adult ChCs (*15*, *17*), while FLP recombinase (FlpO) was driven by the *Pvalb* promoter, a well-established marker for parvalbumin expressing interneurons, including ChCs (*39*). To visualize the ChCs, we crossed the *Vipr2*-*Pvalb* mice with the Ai65 reporter line (*VipR2^Cre^ ^+/-^-Pvalb^flp^ ^+/-^-Ai65^+/-^),* resulting in mice that expressed the tdTomato fluorescent reporter (tdTomato-positive) in cells co-expressing both Cre and FLP recombinases (*38*). Subsequent immunostaining and electrophysiological analysis were performed to verify that the labeled neurons corresponded to ChCs.

The somatic tdTomato labeling was predominantly observed at the boundary of cortical layers L1 and L2/3 (figure 1A), a region densely populated by ChCs (*39*). Notably, tdTomato-positive cells displayed the hallmark features of ChCs (figure 1B, left panel). Using the Filament Tracer tool in Imaris software, we reconstructed their dendritic and axonal architecture (figure 1B, right panel). The axonal arborization revealed the formation of vertical arrays of presynaptic terminals, known as cartridges, while the dendrites extended vertically into L1. These structural characteristics align with previously documented descriptions of ChCs (*10*, *30*). To evaluate the ability of ChCs to target the AIS of PNs, we conducted immunofluorescence staining for the phosphorylated form of IκBα (p-IkBα), a marker enriched in the AIS (*10*, *40*). Surface-surface contact analysis using Imaris algorithms revealed interactions between tdTomato and p-IkBα (a surface contact area of 1672.45 ± 595.03 µm^2^, representing a 13.86 ± 4.81 % out of a total AIS surface area of 9563.64 ± 2127.15 µm^2^, n = 6), confirming that the axonal cartridges of ChCs establish synapses on the AIS of PNs—a distinctive feature of these interneurons (figure 1C-E).

**Figure 1.**
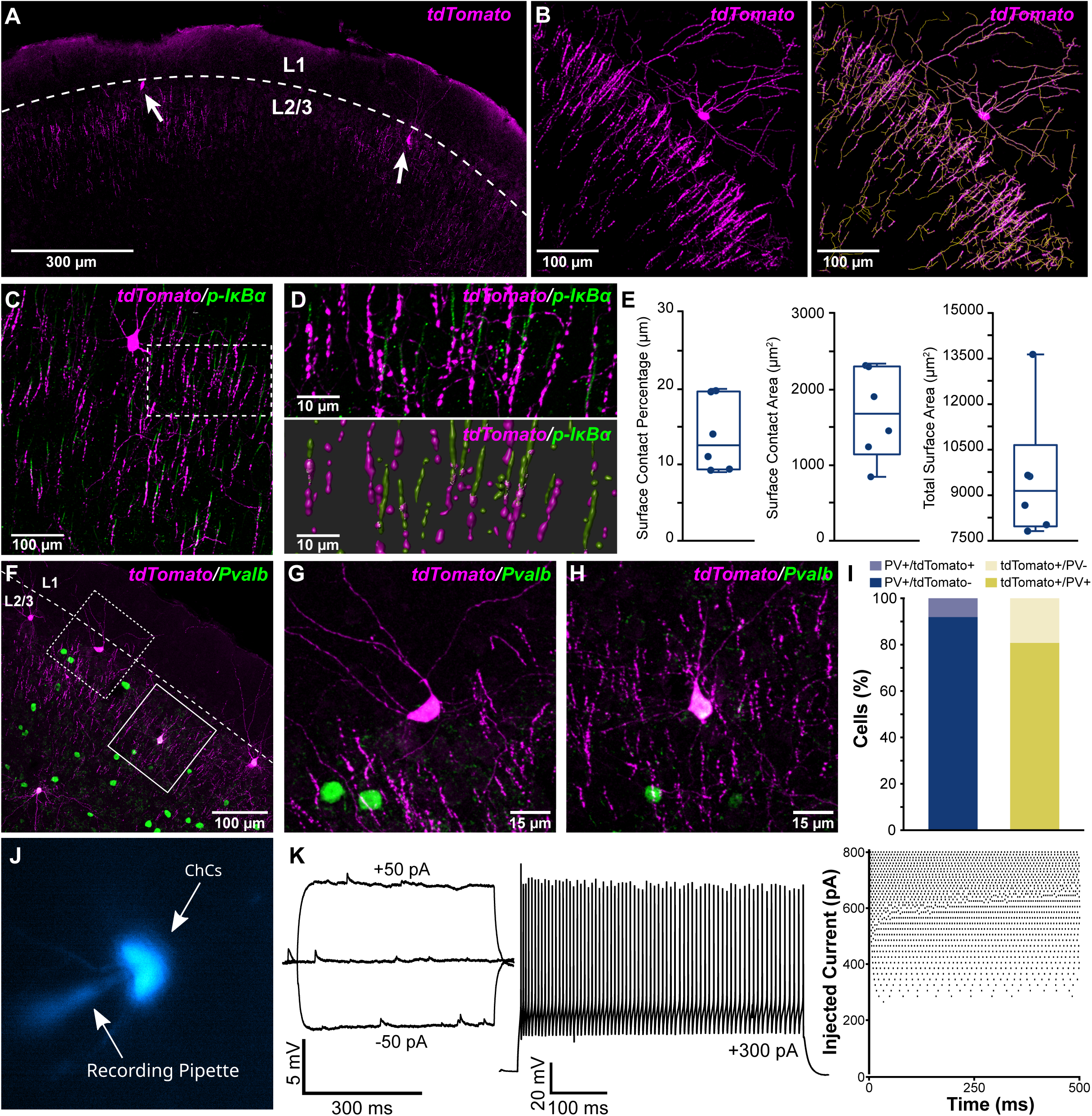
Morphological and electrophysiological identification of Chandelier cells in the secondary motor cortex. **(A)** Representative low-magnification confocal image of a coronal section from the secondary motor cortex. Putative ChCs are identified by their Cre-dependent tdTomato expression (magenta), showing a dense distribution at the L1/L2/3 boundary. White dashed lines indicate the border between L1 and L2/3; arrows point to representative tdTomato-expressing cell bodies. **(B)** Higher-magnification image of a tdTomato-positive ChC in L2/3, with dendrites extending into L1 and axonal cartridges reaching into deeper L2/3 (left). The right panel displays a morphological reconstruction (orange) of the same ChC, generated with the Imaris Filament Tracer tool based on the tdTomato signal (magenta). **(C)** Confocal image of tdTomato (magenta) and phospho-IκBα (p-IκBα, green) immunostaining in L2/3, with the latter marking the axon initial segment (AIS). Note the vertical ‘cartridge’ arrays characteristic of ChCs. Dashed boxes outline regions shown at higher magnification in (D). **(D)** High-magnification images of ChC axonal cartridges (magenta) in close apposition to p-IκBα-labeled AIS segments (green). Detailed views of tdTomato-expressing boutons (magenta) and p-IκBα immunoreactivity (green) reveal their characteristic spatial arrangement. Below, the merged inset shows the superimposed morphological reconstructions of both channels. **(E)** Representative quantifications are shown as box plots for surface contact percentage (left), surface contact area (middle), and total surface area (right) (n = 6 cells, n= 3 mice). **(F)** Representative confocal image illustrating tdTomato-positive cells (magenta) and parvalbumin (green) immunostaining in L1–L2/3. Dashed boxes mark regions enlarged in (G) and (H). **(G, H)** Magnified views of tdTomato-positive cells (magenta) exhibiting the classic ChC morphology, characterized by vertically oriented dendritic trees and axonal cartridges. PV interneurons (green) are visible in the surrounding neuropil. **(I)** Quantification of parvalbumin (PV) and tdTomato co-localization: approximately 90% of PV neurons are tdTomato-negative (blue bars), whereas roughly 80% of tdTomato-positive cells co-express PV (yellow bars). These data indicate that ChCs constitute a minority subpopulation within the broader PV interneuron pool (n= 156 ChCs, n = 4 mice). **(J)** Representative fluorescence-guided patch-clamp recording of a tdTomato-positive ChC (arrowhead) at the L1–L2/3 boundary. The recording pipette is highlighted in pseudocolor blue for visualization. **(K)** Electrophysiological responses recorded from tdTomato-positive neurons. Left: example of a subthreshold membrane response to current injection. Middle: suprathreshold firing in response to a depolarizing current injection. Right: raster plot summarizing the firing patterns across multiple depolarizing current injections. Each dot represents a single action potential event.

Given that ChCs are traditionally classified within the parvalbumin interneuron group, we performed immunofluorescence experiments to assess the degree of colocalization between parvalbumin-expressing neurons and tdTomato-labeled cells (figure 1F-I). Quantification revealed that 93% of parvalbumin positive cells (n=1780) were negative for tdTomato (n=1654), indicating that ChCs represent a minority subgroup within parvalbumin neurons (figure 1I). Additionally, 80.8% of tdTomato-positive cells (n=156) were parvalbumin positive (n=126), supporting their expected ChC identity. Interestingly, 19.2% of tdTomato-positive cells (n=156) did not colocalize with parvalbumin-positive neurons (n=30), suggesting variability in parvalbumin expression among ChCs in the neocortex (figure 1I), consistent with previous findings (*10*). It would indicate that, using our targeting strategy, we may be selecting a subset of ChCs (*VipR2^+^-Pvalb^+^*).

To functionally characterize the putative ChCs, we conducted electrophysiological recordings of tdTomato-positive cells located at the boundaries of cortical layers L1 and L2/3. Fluorescence-guided whole-cell patch-clamp recordings were employed to confirm the neuronal identity (figure 1J). Passive and active membrane properties were measured using current-clamp recordings (figure 1K). On average, tdTomato-positive cells exhibited characteristics consistent with those reported for fast-spiking ChCs (*10*, *41*, *42*), including their firing patterns and intrinsic properties. These included a resting membrane potential of -71.52 ± 0.48 mV, input resistance of 129.95 ± 5.96 MΩ, time constant of 6.51 ± 0.24 ms, rheobase of 206.50 ± 13.09 pA and afterhyperpolarization (AHP) amplitude of 23.22 ± 0.45 mV (n = 32; figure S1). To validate the specificity of our *Vipr2-Pvalb-Ai65* model, it was essential to distinguish ChCs from the broader population of parvalbumin-expressing (PV) interneurons. While *Pvalb-IRES-Cre* mice are commonly used to target PV cells, this line labels a heterogeneous group in which basket cells (BCs) are the predominant, but not the sole, subtype (*7*). To ensure our intersectional approach did not include other PV subpopulations, we compared the electrophysiological profiles of our putative ChCs with those of the general PV population recorded from *PVCre-Ai14* mice. Recordings were conducted on PV cells in layer 2/3 of M2, avoiding the L1/L2/3 boundary region. PV cells (n=30) displayed a stronger fast-spiking phenotype compared to the putative ChCs, and they exhibited distinct electrophysiological parameters (figure S1). These results provide strong evidence that the *VipR2-Pvalb-Ai65D* mouse model is a reliable tool for detailed anatomical and functional analyses of ChCs.

### Direct Modulation of Chandelier Cells by Nicotinic Acetylcholine Receptors

ChCs possess dendrites that extend into L1, suggesting they act as circuit switches by integrating inputs from cortical areas. The dense cholinergic innervation in L1 (*31*) suggests that ChCs could be influenced by cholinergic modulation.

To explore this, we conducted whole-cell patch-clamp recordings in M2 brain slices, targeting ChCs labeled with tdTomato. Using current-clamp configuration, bath application of the cholinergic agonist carbachol (25 µM) induced a reversible depolarization of the ChCs resting membrane potential (8.24 ± 0.22 mV, n = 10) (figure S2A). Similar results were observed with the nicotinic agonist DMPP (25 µM), suggesting the involvement of nAChRs (10.18 ± 0.27 mV, n = 17) (figure S2B). Remarkably, the depolarizing response to DMPP persisted despite the application of a blocking solution containing TTX (1 µM), CNQX (10 µM), DL-APV (50 µM), picrotoxin (50 µM), and atropine (4 µM), confirming that the effect was independent of synaptic transmission and specific for nicotinic modulation (figure S2C) (6.87 ± 0.60 mV, n = 20). To confirm that the depolarization of ChCs was primarily mediated by the activation of nAChRs, we performed voltage-clamp recordings while applying DMPP via local pressure (100 µM, 20 ms, 4 psi; figure 2A). This application elicited an inward current in normal ACSF (figure 2B; amplitude: -76.38 ± 7.89 pA, n = 23) that persisted in the presence of the previously described blocking solution (amplitude: -47.37 ± 4.69 pA, n = 23) (figure 2B). Notably, the inward current was abolished upon the addition of mecamylamine (MMA, 5 µM), a specific nAChR antagonist (amplitude: N/A, n = 23) (figure 2B). These findings confirm the key role of nAChRs in mediating the observed response.

**Figure 2.**
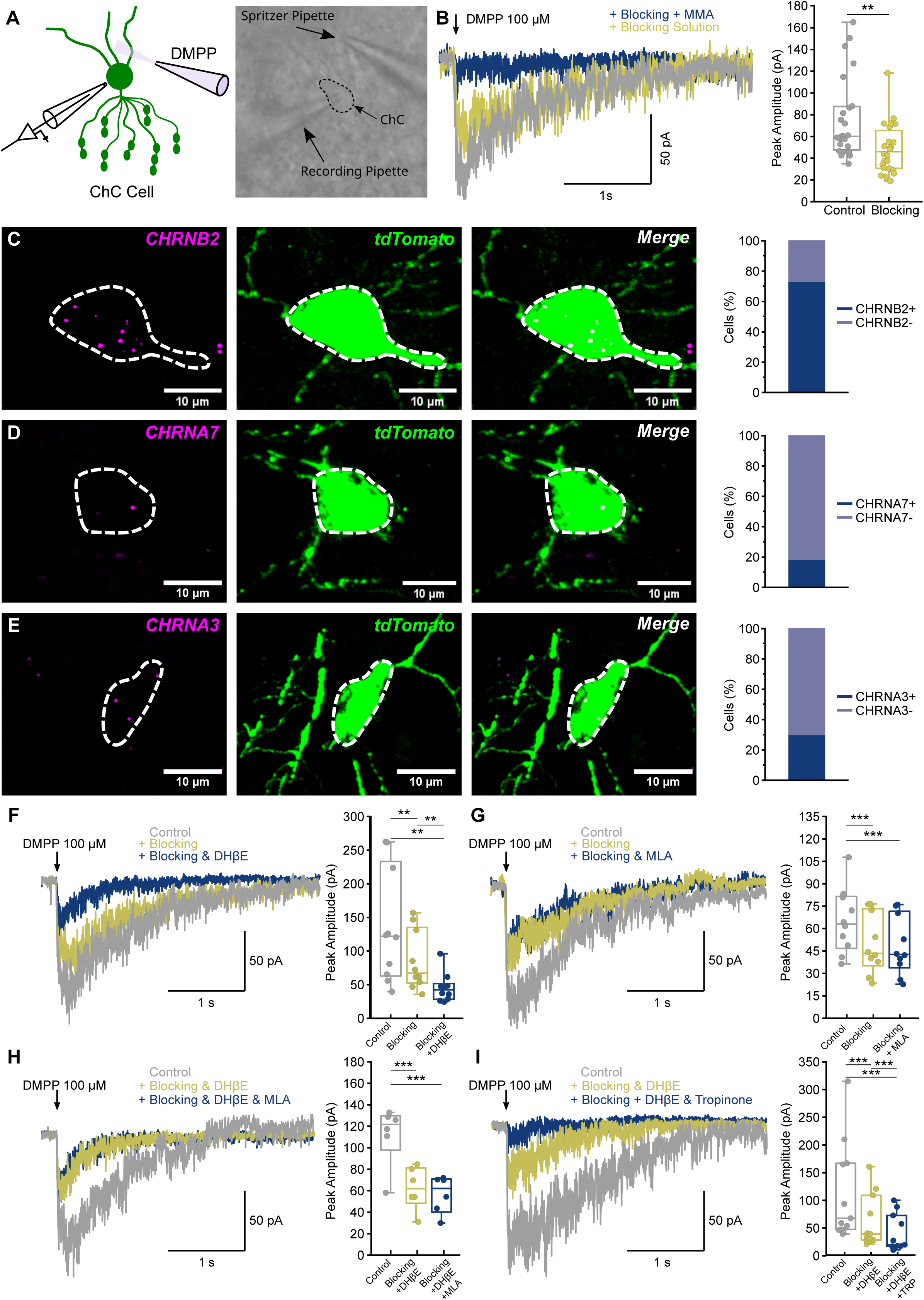
Heteromeric nicotinic acetylcholine receptors mediate cholinergic excitation in cortical Chandelier Cells. **(A)** Left, schematic of the experimental setup illustrating a tdTomato-positive ChC targeted for whole-cell patch-clamp recording during focal pressure ejection (puff) of the nicotinic agonist DMPP. Right, representative infrared (IR-DIC) micrograph showing the ChC approached by both the recording electrode and the drug delivery pipette. **(B)** Left, Representative whole-cell patch-clamp recordings showing the contribution of nAChRs to DMPP-evoked inward currents in ChCs. The grey trace represents the response under control conditions; the yellow trace shows the response during broad synaptic and muscarinic blockade (TTX, 1µM; atropine, 4µM; CNQX, 10 µM; picrotoxin, 50 µM; AP-V, 50 µM). The blue trace illustrates the inhibition of the current following the subsequent application of mecamylamine (MMA, 5 µM), a non-selective nAChR antagonist. Right, box plots quantifying the DMPP-evoked inward currents under control and blocking conditions. Data for mecamylamine are not shown in the boxplots because the drug-induced blockade was complete, reducing the inward currents to baseline levels and preventing reliable quantification; (p < 0.01, Wilcoxon signed-rank test, n=23 cells, 11 mice). **(C–E)** Fluorescent *in situ* hybridization (FISH) of nicotinic acetylcholine receptor (nAChR) subunits in tdTomato-labeled ChCs. White dashed lines indicate the boundaries of the sampled somata. (C) *Chrnb2* mRNA (magenta) is detected in the majority of ChCs (73.78%). (D) *Chrna7* mRNA (magenta) is present in a smaller subset (17.86%). (E) *Chrna3* mRNA (magenta) is observed in approximately 30% of ChCs. **(F–I)** Right, representative whole-cell patch-clamp recordings showing subunit-specific contributions to DMPP-evoked inward currents in ChCs. Grey traces represent responses under control conditions; yellow traces show responses during broad synaptic and muscarinic blockade (TTX, 1 µM; atropine, 4 µM; CNQX, 10 µM; picrotoxin, 50 µM; AP-V, 50 µM). Blue traces illustrate the remaining current following the subsequent application of specific nicotinic antagonists: (F) the β2-selective antagonist DHβE (10 µM), (G) the α7-selective antagonist methyllycaconitine (MLA, 200 nM), (H) sequential application of DHβE and MLA, and (I) the α3-selective antagonist tropinone (300 µM) following DHβE. Right, box plots summarize the peak amplitudes across experimental conditions. (**p < 0.01, *p < 0.001; Friedman test followed by Wilcoxon signed-rank test with Bonferroni correction for F, H, I; one-way ANOVA with Tukey’s post-hoc test for G, n=10 cells, 8 mice (F), n=10 cells, 9 mice (G), n=6 cells, 3 mice (H), n=11 cells, 9 mice. (I).

### Heteromeric Nicotinic Receptors Mediate the Cholinergic Response in Chandelier Cells

Previous studies, including transcriptomic analyses, have reported the presence of cholinergic receptor subunits in ChCs (*38*, *43*, *44*). However, their precise subunit composition, subcellular localization, and the specific modes of transmission in which they participate remain incompletely defined. To address this gap, we performed fluorescent in situ hybridization (FISH) (figure 2C-E). To enhance the number of ChCs available for this quantification, we modified our experimental approach slightly. To overcome the low labeling density observed in the *Vipr2-Pvalb-Ai65* line, we implemented an optimized strategy by stereotaxically injecting a Cre-dependent tdTomato AAV (AAV9-hSynP-Flex-tdTomato) into the M2 region of adult *Vipr2-Cre* mice (*15*, *17*). This approach exploits the adult-specific expression of the *Vipr2* promoter in ChCs, resulting in a significantly higher number of labeled cells compared to our initial triple-transgenic model. Three weeks after the injection, the FISH experiments were performed. The majority of ChCs (73.78%, n = 166/229) expressed the β2 nicotinic receptor subunit (CHRNB2), with a mean fluorescence intensity of 29.82 ± 0.9 a.u (figure 2C). A smaller subset of ChCs expressed the α7 subunit (CHRNA7, 17.86%, n = 20/112), also with a mean fluorescence intensity of 15.88 ± 1.55 a.u (figure 2D). These findings suggest that heteromeric nAChRs are the predominant subtype in ChCs. To further dissect the contributions of specific nAChR subunits, we performed patch-clamp recordings using the protocol described in figure 2A but applying subunit-specific blockers instead of a general nAChR antagonist (figure 2F-H). To assess the role of the β2 subunit, DMPP application under control conditions evoked an inward current with a peak amplitude of -125.83 ± 25.99 pA (n = 10). This response was reduced to -78.96 ± 13.94 pA (n = 10) following the blockage of synaptic transmission and further attenuated to -2.04 ± 6.91 pA (n = 10) upon the addition of DHβE (10 µM), a selective antagonist of β2-containing nAChRs, to the ACSF (figure 2F).

To investigate the role of the α7 subunit, we applied DMPP under control conditions, which elicited an inward current with an amplitude of -64.97 ± 6.86 pA (n = 10). This response was reduced after blocking synaptic transmission (-49.38 ± 6.21 pA; n = 10) but remained unaffected by the addition of methyllycaconitine (MLA), a selective antagonist of α7-containing nAChRs (200 nM, -48.39 ± 6.18 pA, p > 0.05, n = 10; figure 2G). Given that α7-containing nAChRs are characterized by their rapid inward current kinetics (*45*), potentially overshadowed by the dominant β2-containing nAChRs, we further examined the inward current by sequentially applying DHβE and MLA. Under control conditions, the amplitude was -112.09 ± 11.29 pA (n = 6), which decreased to -62.19 ± 8.11 pA (n = 6) after blocking synaptic transmission in the presence of DHβE and to –56.61 ± 7.04 pA (n = 6) following the addition of MLA. The lack of a significant difference between the DHβE and DHβE + MLA conditions (p > 0.05; figure 2H) indicates that α7-containing nAChRs have a limited role in the cholinergic modulation of ChCs. This result underscores the predominant involvement of β2-containing nAChRs in mediating the cholinergic regulation of ChCs.

Since a residual inward current persisted after blocking with MLA and DHβE, we hypothesized that additional heteromeric receptor subtypes might contribute to the nicotinic modulation of ChCs. Transcriptomic studies in the mouse cortex have previously reported the expression of the α3 subunit (CHRNA3) (*38*, *46*). To investigate the potential involvement of α3 in heteromeric nAChRs, we performed fluorescent in situ hybridization, which confirmed the presence of α3 subunit in nearly 30% of ChCs (CHRNA3, 29.52%, n = 31 out of 105) with a mean fluorescence intensity of 36.57 ± 3.25 a.u (figure 2E). To assess the contribution of α3 subunits to the inward current, we applied tropinone, a specific α3 subunit blocker. Under control conditions, the inward current amplitude was -122.24 ± 28.32 pA (n = 10). This amplitude was reduced to -65.74 ± 15.32 pA (n = 10) after blocking synaptic transmission in the presence of DHβE and further decreased to -42.90 ± 10.62 pA (n = 10) when tropinone (300 µM) was added alongside DHβE (figure 2I).

Collectively, these findings highlight the involvement of heteromeric nAChRs, including those containing the α3 subunit, in the cholinergic modulation of ChCs.

### Basal forebrain cholinergic activation evokes slow, sustained nicotinic responses in Chandelier cells

The principal cholinergic nuclei that project to the neocortex are located in the BF. To assess whether these neurons drive nicotinic modulation of ChCs, we combined optogenetic activation of BF cholinergic axons with whole-cell patch-clamp recordings in acute cortical slices.

We used a transgenic mouse line enabling simultaneous identification of cortical ChCs and Cre-dependent targeting of cholinergic neurons (*VipR2^Cre^ ^+/-^-Pvalb^flp^ ^+/-^-Ai65^+/-^-ChAT^Cre+/-^*). To selectively activate cholinergic projections, we injected a Cre-dependent AAV encoding ChR2-YFP into the primary cholinergic nuclei (figure 3A–C). Three weeks after injection, immunofluorescence analyses confirmed ChR2-expressing cholinergic axons within the M2 cortical region (figure 3D).

**Figure 3.**
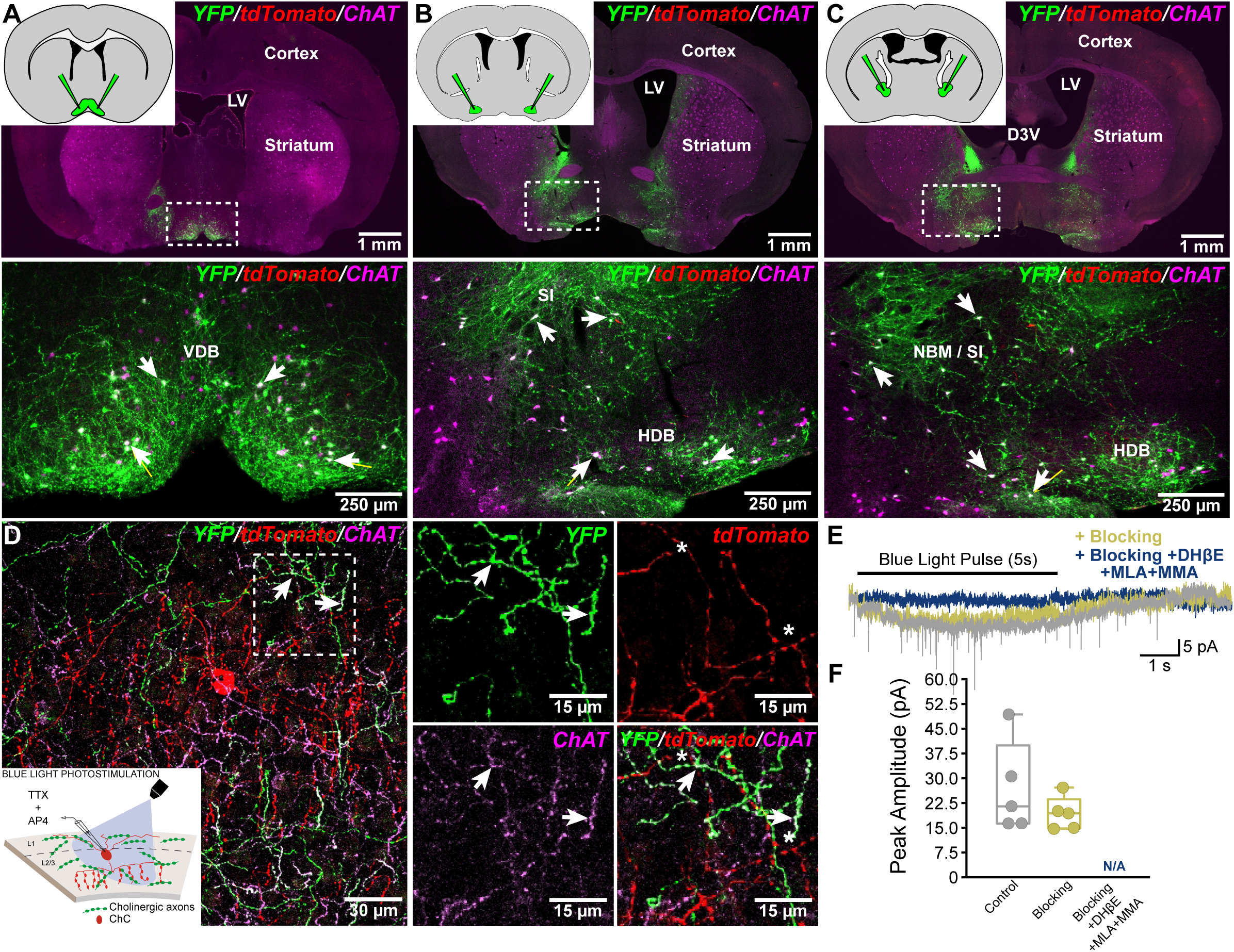
Optogenetic activation of basal forebrain cholinergic inputs drives nicotinic currents in cortical Chandelier Cells. **(A–C)** Top panels, schematic insets indicate three rostro-caudal levels of the injection sites (see Methods for coordinates). Panoramic images show ChR2-YFP (green) and tdTomato-positive ChCs (red) overlaid with choline acetyltransferase (ChAT) immunostaining (magenta). Major forebrain structures, including the cortex, striatum, and lateral ventricles (LV), are outlined. Boxed regions are enlarged in the panels below. Bottom panels, representative high-magnification images of the vertical diagonal band (VDB), horizontal diagonal band (HDB), and nucleus basalis/substantia innominata (NBM/SI) reveal dense cholinergic labeling. White arrows highlight examples of YFP-positive/ChAT-positive co-labeled neurons **(D)** High-magnification confocal images of cortical cholinergic terminals in close proximity to ChC dendrites. Left, merged image showing YFP-positive (green) and ChAT-positive (magenta) cholinergic axons interspersed with tdTomato-labeled ChC dendrites (red) in L2/3. The boxed area is shown at higher resolution on the right, where white arrows indicate cholinergic varicosities (co-labeled for YFP and ChAT) in close apposition to ChC dendritic segments. The inset schematizes the **(E)** Whole-cell patch-clamp recordings showing light-evoked postsynaptic currents. Grey traces represent responses under control conditions; yellow traces show responses during broad synaptic and muscarinic blockade (TTX, 1 µM; atropine, 4 µM; CNQX, 10 µM; picrotoxin, 50 µM; AP-V, 50 µM). Blue traces illustrate the inhibition of the remaining nicotinic current following the subsequent application of specific nAChR antagonists: DHβE (10 µM), methyllycaconitine (MLA, 200 nM), and mecamylamine (MMA, 5 µM) **(F)** Box plots summarizing the peak amplitudes of light-evoked currents across the experimental conditions described in (E), n = 5 cells, 2 mice.

We then performed patch-clamp recordings from visually identified ChCs while optogenetically stimulating cholinergic afferents in M2 (figure 3D, inset). Brief photostimulation (2, 5 or 10 ms pulses) failed to evoke measurable responses, whereas prolonged photostimulation (5 s) elicited a robust inward current in ChCs under both ACSF and the previously described synaptic-blocking conditions. This current was completely abolished by nicotinic antagonists (MMA, 5 µM; MLA, 200 nM; DHβE, 10 µM; n = 5; figure 3E–F). The mean current amplitude was -26.76 ± 6.19 pA under control conditions and -19.21 ± 2.26 pA in the blocking solution; no response was detectable after nicotinic receptor blockade.

These findings demonstrate that BF-derived cholinergic fibers drive nicotinic activation of ChCs. The long latency, gradual onset, and sustained plateau of the optogenetically evoked currents are not characteristic of fast, point-to-point synaptic transmission. Instead, their temporal profile is consistent with a spatially diffuse mode of cholinergic signaling, potentially involving extracellular acetylcholine acting on high-affinity heteromeric nAChRs distributed along ChC somatodendritic compartments.

### Sustained Elevation of Extracellular Acetylcholine Enhances Intrinsic Excitability of Chandelier Cells

If cholinergic signaling onto ChCs operates through a temporally extended and potentially diffuse mechanism, then sustained elevations in extracellular acetylcholine should be sufficient to modulate their intrinsic excitability. To directly test this possibility, we examined the effects of acetylcholinesterase inhibition using physostigmine, thereby increasing ambient acetylcholine levels in the slice preparation.

Whole-cell patch-clamp recordings were performed in acute cortical slices under pharmacological blockade of fast synaptic transmission (CNQX, 10 µM; DL-APV, 50 µM; and picrotoxin, 50 µM) and muscarinic receptors (atropine, 4 µM). After characterizing baseline intrinsic properties, bath application of physostigmine (10 µM) induced gradual and sustained enhancements in excitability (n = 12; figure 4A-C). These alterations included a significant depolarization of the resting membrane potential (from -66.38 ± 0.72 mV to -62.93 ± 1.29 mV; figure 4A), a reduction in rheobase (from 143.5 ± 12.02 pA to 120.4 ± 9.56 pA; figure 4B), and a prominent leftward shift in frequency–current (F–I) curves (Figure 4C). These effects developed over several minutes and persisted throughout drug application, consistent with the progressive accumulation of endogenous acetylcholine. To confirm the nicotinic component involved, a separate cohort of experiments was conducted in the presence of nicotinic antagonists (MLA, 200 nM; MMA, 100 µM) (n = 10; figure 4D-F). Under these conditions, the physostigmine-induced depolarization of resting membrane potential was completely abolished (-69.05 ± 1.02 mV vs. -68.24 ± 1.16 mV; figure 4D) as was the shift in rheobase (170 ± 20.71 pA vs. 164.6 ± 20.54 pA; figure 4E), with no change in firing frequency (figure 4F). These results indicate that the enhancement of neuronal output and the shift in resting state are strictly mediated by nicotinic receptor activation.

**Figure 4.**
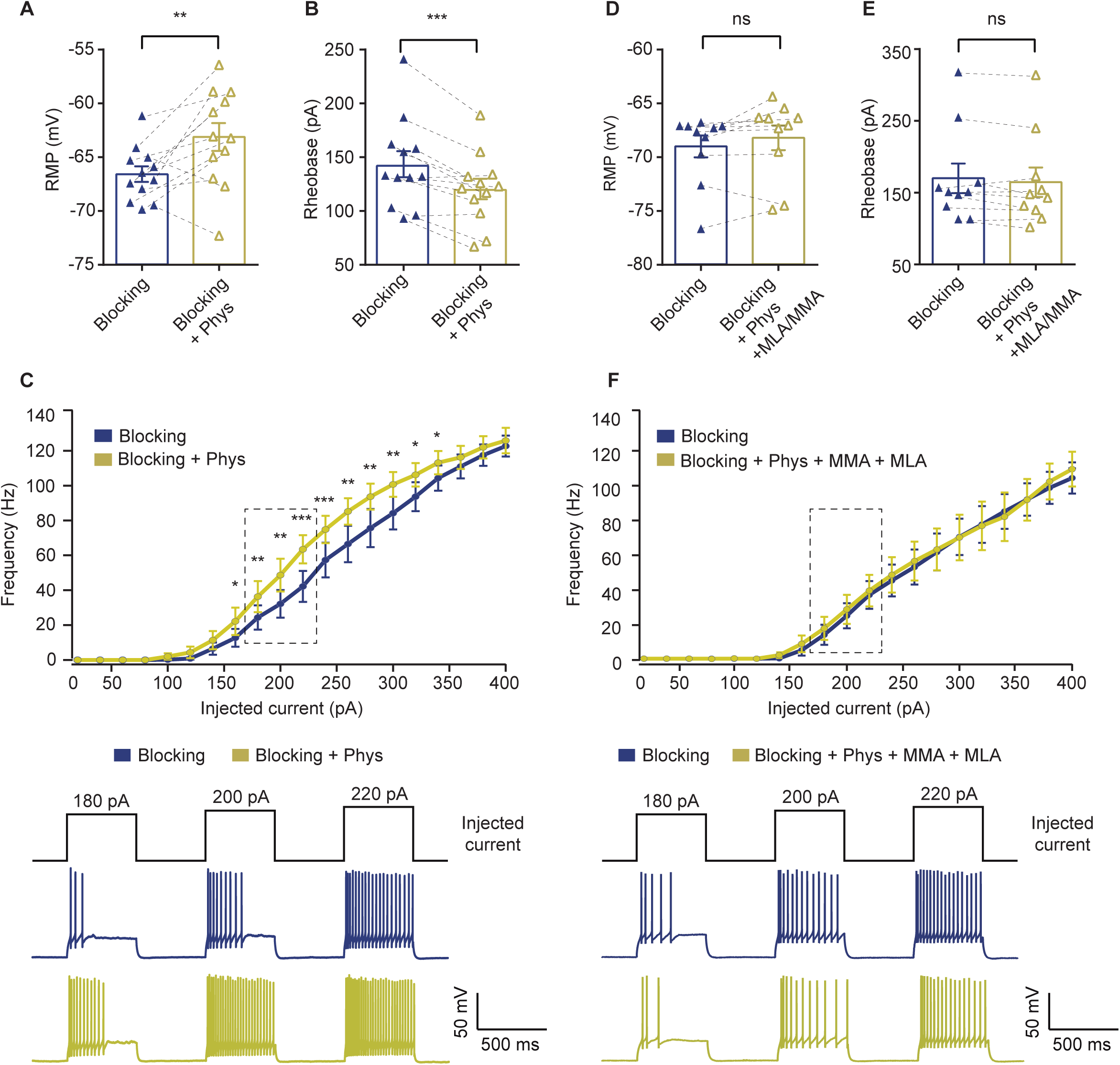
Nicotinic receptor blockade abolishes physostigmine-induced enhancement of Chandelier Cell excitability. **(A,B)** Resting membrane potential and rheobase under baseline blocking conditions and after bath application of physostigmine (10 µM). Baseline recordings were performed in the presence of synaptic blockers (CNQX, 10 µM; DL-APV, 50 µM; picrotoxin, 50 µM) and atropine (4 µM). Physostigmine significantly depolarized the membrane (**p < 0.01, paired t-test; n = 12 cells from 3 mice) and significantly lowered rheobase (***p < 0.001, paired t-test; n = 12 cells from 3 mice). **(C)** Frequency–current (F-I) relationships comparing blocking conditions with blocking + physostigmine. Physostigmine increased firing across a range of injected currents (*p < 0.05, **p < 0.01, ***p < 0.001; paired t-tests, n = 12 cells, 3 mice). The dashed box indicates the range of current injections from which the representative traces shown below were selected. **(D,E)** RMP and rheobase measured under blocking conditions and after physostigmine in the presence of nicotinic receptor antagonists methyllycaconitine (MLA, 200 nM) and mecamylamine (MMA, 100 µM). Under nicotinic blockade, physostigmine did not significantly alter resting membrane potential or rheobase (ns, paired t-test; n = 10 cells from 3 mice for both measurements). **(F)** Frequency–current (F-I) relationships comparing blocking conditions with blocking + physostigmine + MMA + MLA. Nicotinic receptor blockade prevented the physostigmine-induced increase in excitability. The dashed box indicates the range of current injections from which the representative traces shown below were selected.

### Chandelier Cell Activity Correlates with High Arousal States in the M2 Cortex

Fluctuations in arousal states profoundly impact cortical activity and behavioral performance. ACh plays a pivotal role in these dynamics, modulating the activity of individual cortical neurons and orchestrating their interactions within neural networks. Cholinergic signaling is strongly linked to high-arousal states, which often accompany active behaviors such as locomotion, an observable measure of internal arousal levels, and spontaneous pupil diameter fluctuations that reflect shifts in vigilance, attention, and cognitive effort (*35*, *47–49*).

To explore the relationship between ChCs activity and behavioral states, we utilized two-photon calcium imaging to monitor ChCs activity in awake, head-fixed mice navigating on a floating spherical treadmill. Behavioral metrics, including pupil diameter and running speed, were simultaneously recorded during imaging. To achieve this, we injected an adeno-associated viral vector encoding a CRE-dependent calcium indicator gene, GCaMP6s, into the M2 cortical region of adult *VipR2^Cre^ ^+/-^-Pvalb^flp^ ^+/-^-Ai65^+/-^* mice (figure 5A). Since CRE recombinase is expressed in these mice under the control of the *Vipr2* promoter, we observed GCaMP6s expression in both tdTomato-positive (30.5%) and tdTomato-negative (69.5%) cells. To determine whether tdTomato-negative cells displayed ChC characteristics, we performed immunofluorescence against GCaMP6s, identifying the typical ChC morphology (figure 5B), consistent with previous studies (*17*). Additionally, 72.1% of GCaMP6s-positive cells were also positive for parvalbumin, while 27.9% were negative, further suggesting variability in parvalbumin expression among ChCs (3 mice and 15 slices; figure 5C).

**Figure 5.**
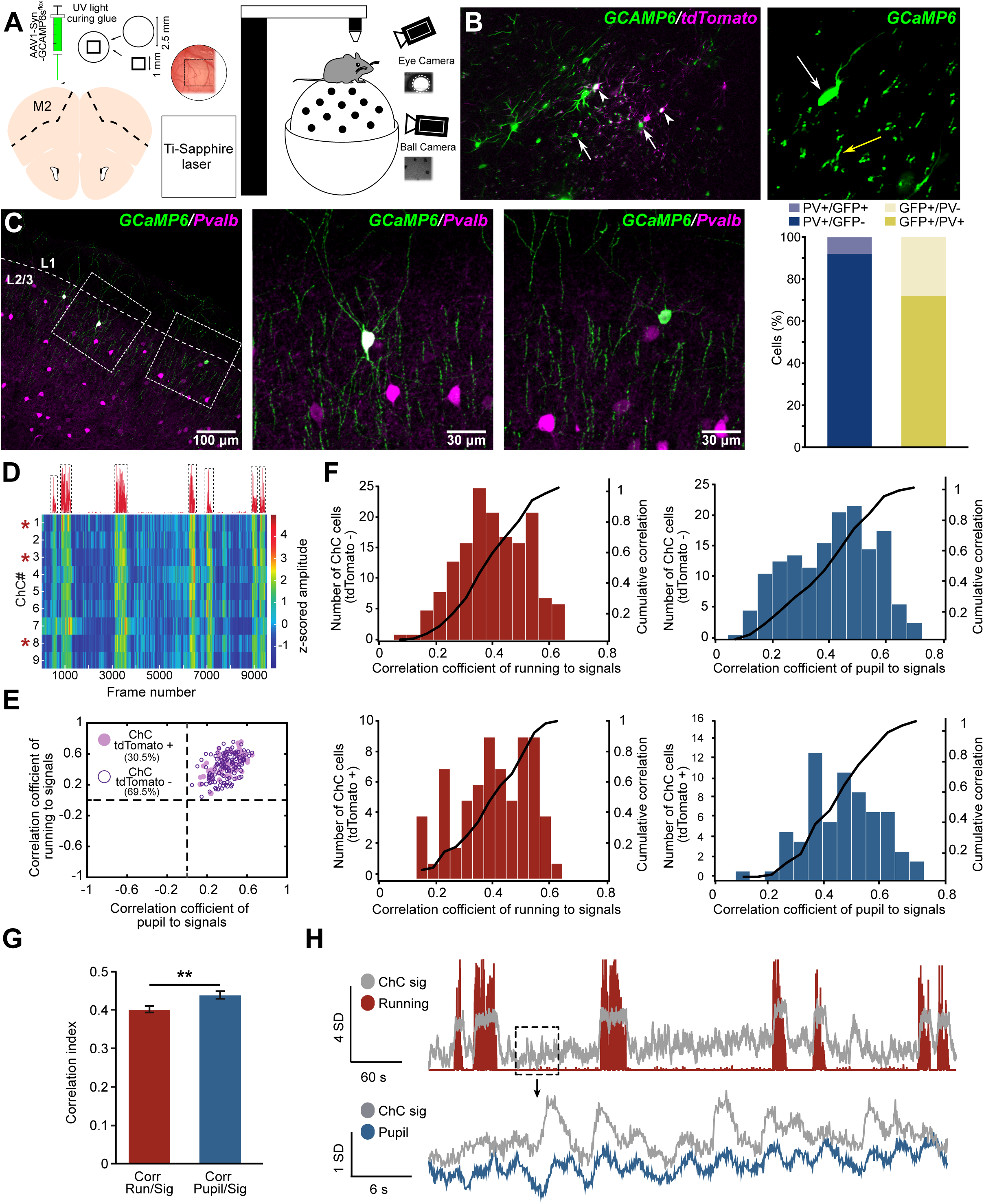
Chandelier Cells in the secondary motor cortex are robustly activated by locomotion and arousal. **(A)** Schematic overview of the experimental approach. Adult *VipR2-Cre+/-; PvalbFlp+/-; Ai65+/-* mice were injected in the secondary motor cortex (M2) with a Cre-dependent AAV expressing GCaMP6s. During two-photon calcium imaging, head-fixed mice navigated on a floating spherical treadmill, and pupil diameter and running speed were recorded simultaneously **(B)** Representative two-photon images from layer 2/3 of the M2 cortex showing the expression of GCaMP6s (green) and tdTomato (magenta). Arrowheads indicate dual-labeled ChCs expressing both GCaMP6s and tdTomato, while white arrows point to neurons expressing GCaMP6s alone. Right, high-magnification inset of a single GCaMP6s-labeled ChC, highlighting the soma (white arrow) and its characteristic axonal cartridges (yellow arrow). **(C)** Immunofluorescence-based confirmation of ChC identity. Left, GCaMP6s (green) labels neurons with distinctive ChC axonal arborizations and cartridge morphology. Parvalbumin (magenta) identifies parvalbumin-positive interneurons. *Middle and right,* higher magnification images of selected areas confirm ChC morphology. *Far right,* stacked bar graphs quantifying cell identity. Left graph, proportion of PV-labelled cells that are GCaMP6s-positive (light blue; n = 111) versus GCaMP6s-negative (dark blue; n = 1305). Right graph, proportion of GCaMP6s-labelled cells that are PV-positive (dark yellow; n = 111) versus PV-negative (light yellow; n = 43). **(D)** Representative activity heatmap illustrating calcium transients for individual ChCs within a single focal plane over time. The top red trace indicates the animal’s locomotor activity during the corresponding imaging session. Red arrowheads denote dual-labeled ChCs (tdTomato-positive/GCaMP6s-positive). Note that the temporal activity patterns are consistent between GCaMP6s-expressing neurons, regardless of tdTomato expression. **(E)** Scatter plot illustrating the Pearson’s correlation coefficients for individual ChCs. Each data point represents the correlation of a single cell’s calcium activity relative to pupil dilation (x-axis) and ball movement (y-axis). Filled circles denote tdTomato-positive ChCs, while open circles represent tdTomato-negative neurons (n = 226 cells, 3 mice). Note that all cells cluster in the upper-right quadrant, indicating a consistent positive correlation with both locomotion and arousal levels, regardless of tdTomato expression. **(F)** Histograms of Pearson’s correlation coefficients for locomotor activity (red) and pupil diameter (blue). The distributions for tdTomato-positive ChCs (n = 69 cells) and tdTomato-negative neurons (n = 157 cells) indicate that both populations exhibit similar coupling to behavioral state and arousal. (n.s., p > 0.05, Kolmogorov-Smirnov test). **(G)** Bar plot showing the mean correlation between ChC activity and either locomotion (red) or pupil diameter (blue). Data from both tdTomato-positive and tdTomato-negative populations were pooled, as they showed no significant functional differences. ChC activity is significantly correlated with both behavioral parameters. Data are presented as mean ± SEM. (p < 0.01, Mann-Whitney U test, n = 226 cells, 3 mice). **(H)** Top, representative time series from a single imaging session demonstrating that ChC calcium transients (grey) closely track locomotor bouts (red trace). Bottom, magnified view of the boxed region from the upper panel, illustrating that the calcium signal follows pupil fluctuations even during resting periods, in the absence of locomotion.

Qualitative analysis of calcium signals from both tdTomato-positive and tdTomato-negative ChCs revealed increased activity during periods of treadmill running (figure 5D). To quantify the association between arousal state and ChC responses, we calculated correlation coefficients between the activity of both tdTomato-positive and tdTomato-negative ChCs and two key behavioral metrics: pupil size and running speed. Our analysis showed that all imaged neurons (tdTomato-positive and tdTomato-negative) exhibited positive correlations with both pupil diameter and running speed (figure 5E and video 1). Since the degree of correlation did not significantly differ between the two populations, we combined them for subsequent analyses.

The correlation analysis between ChCs activity and individual behavioral metrics, such as running and pupil diameter, revealed a significantly stronger relationship with pupil size than with locomotion (figure 5F-G). This finding likely reflects that, during quiet periods, behavioral changes are not detected as overt movement but are instead linked to spontaneous fluctuations in pupil diameter (figure 5H). These fluctuations closely align with ChC activity, offering a more sensitive indicator of underlying neural dynamics. Previous studies have shown that pupil dynamics predict changes in oscillatory network activity and are reliably correlated with subthreshold membrane potential fluctuations in certain cortical neuron subtypes, even in the absence of overt movement (*33*). These membrane potential changes result from alterations in cholinergic and adrenergic inputs (*50*). Interestingly, pupil dynamics emerge as a robust proxy for tracking neuromodulatory changes within the cortex (*35*). Given that we have demonstrated ChCs are modulated by cholinergic input, it is plausible to hypothesize that this neuromodulation has an important role in ChC function.

### Cholinergic Blockade Attenuates Chandelier Cells Neuronal Responses to Locomotion

Cholinergic modulation plays a key role in shaping the activity of cortical interneurons during behaviorally relevant states such as locomotion (*34*, *50*). Because ChCs are modulated by nAChRs, we checked how the blockade of cholinergic signaling impacts ChC responses during locomotion.

We performed in vivo two-photon calcium imaging of the same ChCs in the presence and absence of nicotinic antagonists (MMA, 1 mM; MLA, 0.1 mM) (*34*). Blocking nicotinic receptors led to a marked reduction in ChC responses during locomotion (figure 6A). Cross-correlation analysis between calcium signals and treadmill movement showed a sharp decline after antagonist treatment (Zero-time cross-correlation: Control: 0.33 ± 0.03; MMA/MLA: 0.08 ± 0.03, n = 20) (figure 6B). Additionally, quantitative analysis revealed a significant decrease in the normalized amplitudes of calcium signals associated with movement following antagonist treatment (Control: 1.41 ± 0.17 SEM; MMA/MLA: 0.05 ± 0.13 SEM, n = 20) (figure 6C). These findings highlight the critical role of cholinergic inputs in modulating ChC activity and ensuring their proper tuning during active behavioral states.

**Figure 6.**
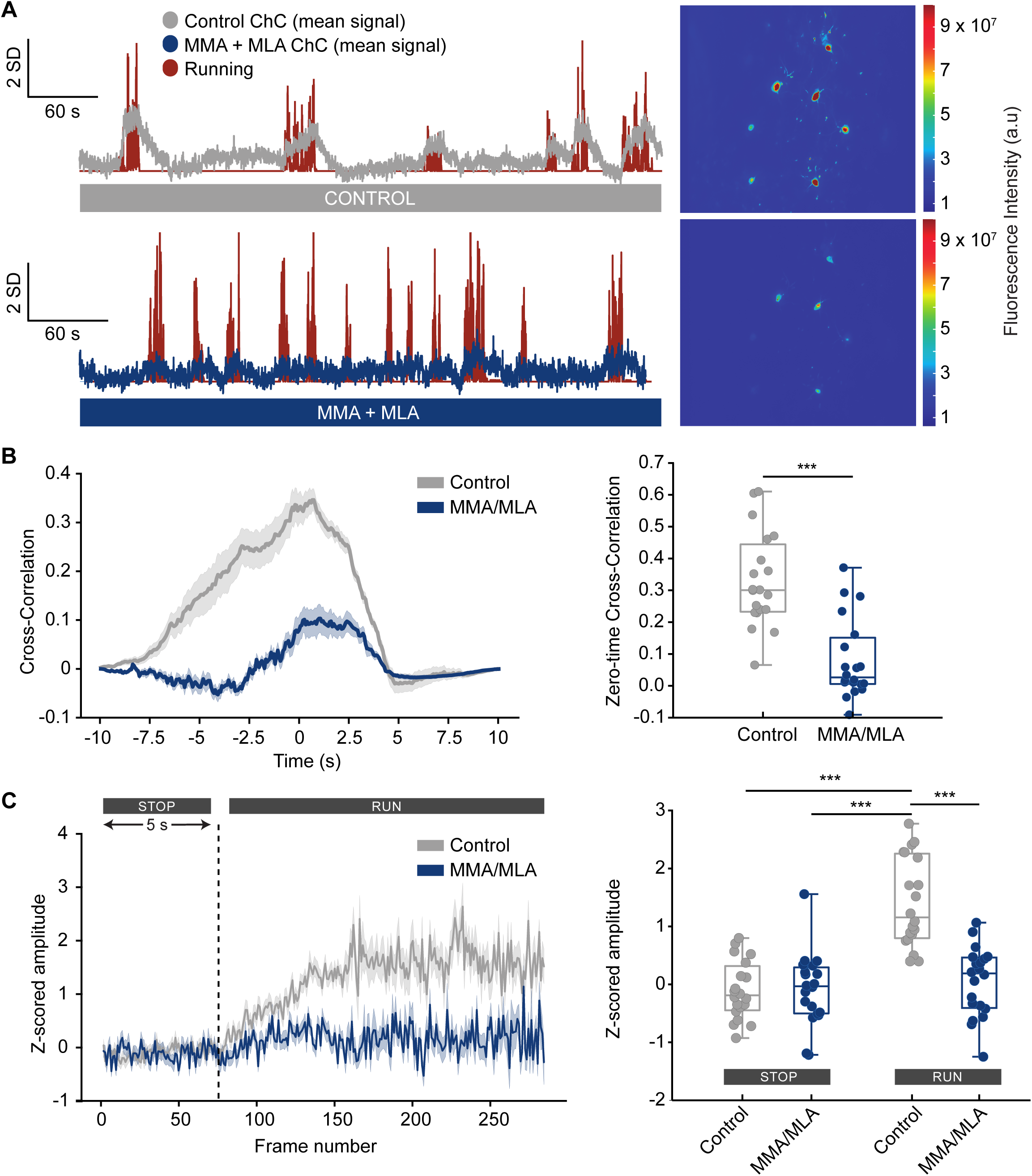
Nicotinic signaling is required for the behavioral modulation of neocortical Chandelier Cells. **(A)** Representative calcium imaging traces showing the mean z-scored activity of ChCs aligned to locomotor bouts (red trace). Top, under control conditions, z-scored calcium signals (grey trace) are closely time-locked to running behavior. Bottom, following bath application of the nAChR antagonists mecamylamine (MMA) and methyllycaconitine (MLA), the locomotion-induced z-score fluctuations are markedly suppressed (blue trace). Right, corresponding pseudocolor images illustrating the standard deviation of GCaMP6s fluorescence within the field of view. The bottom image demonstrates a significant reduction in activity-dependent fluorescence fluctuations following nAChR blockade compared to control (top). **(B)** Left, cross-correlation analysis between ChC calcium transients and locomotor activity. The plot shows the mean cross-correlation over time in control conditions (grey) and after local perfusion of the nAChR antagonists MMA and MLA (blue). In control conditions, a prominent central peak near zero-lag indicates high synchrony between neuronal activity and movement, which is markedly reduced upon nicotinic receptor blockade. Data are presented as mean ± SEM (shaded areas). Right, boxplots depicting the distribution of cross-correlation values at zero lag (time = 0 s) for control and MMA/MLA conditions. Zero-lag cross-correlation is significantly reduced in the presence of MMA/MLA compared to control (*p* < 0.001, Wilcoxon signed-rank test, n = 20 cells). **(C)** Left, average running-associated calcium transients under control (grey) and MMA/MLA (blue) conditions. Signals are aligned to the onset of locomotion (vertical dashed line). The grey trace shows a clear increase in z-scored amplitude upon running, which is significantly blunted following nicotinic receptor blockade (blue trace). Data are presented as mean ± SEM (shaded areas). Right, box plots quantifying the mean z-scored amplitudes during stationary (’STOP’) and locomotor (’RUN’) states. Under control conditions, ChC activity significantly increases during running, an effect that is markedly abolished by the application of MMA/MLA. (*p < 0.001, one-way ANOVA with Tukey’s post-hoc test, n = 20 cells).

## Discussion

Our findings reveal that ChCs in the M2 cortical region are strongly modulated by cholinergic signaling through nAChRs, with this modulation closely tied to arousal states and supported by a mode of action compatible with volume-mediated ACh release.

In this study, we used a genetic intersectional strategy to selectively target and visualize a subpopulation of parvalbumin-expressing ChCs in adult mice by combining *VipR2*- and *Pvalb*-driven recombinases with the *Ai65D* fluorescent reporter line (*38*). Our immunohistochemical analyses indicated that approximately 80% of tdTomato-positive cells co-express parvalbumin, supporting their expected ChC identity, whereas the remaining 20% did not exhibit strong parvalbumin signals (figure 1I), underscoring variability in parvalbumin expression among ChCs in the neocortex—an observation consistent with previous reports (*10*). These results suggest that, while our strategy effectively labels a substantial fraction of ChCs, it likely targets a specific subset of *Vipr2-Pvalb* ChCs and potentially underestimates the overall ChC population. Indeed, recent transcriptomic studies further subdivided the *Vipr2-Pvalb*-expressing ChC population into two main clusters with distinct transcriptomic profiles (*51*), emphasizing the complexity of interneuron classification in the cortex and highlighting the need for increasingly precise genetic tools to dissect the diverse cellular phenotypes and functions of cortical inhibitory interneurons.

Ex vivo patch-clamp recordings confirmed that ChCs are fast-spiking interneurons, characterized by brief action potentials and high-frequency firing comparable to other PV-expressing interneurons (figure S1A-D). We assume in this discussion that the vast majority of parvalbumin-positive interneurons expressing tdTomato in the *PVCre-Ai14* mouse model correspond to BCs, which constitute the population recorded in our electrophysiological experiments. Compared to BCs, ChCs exhibited higher membrane capacitance and a markedly lower maximal firing rate (∼55% of BCs cells), together with a more depolarized AP threshold, slower rise time, and broader half-width, while RMP, input resistance, AHP amplitude, and rheobase were similar (figure S1E). These findings match previous mouse data (*10*), although other studies reported different input resistance, rheobase, and threshold relationships between these cell types (*41*). Cross-species work in rats and monkeys further indicates that ChC–BC differences can invert, with ChCs firing faster and having more negative thresholds than BCs (*52*). Overall, these studies, together with our data, point to substantial electrophysiological heterogeneity of ChCs across species and cortical regions, complicating a unified functional interpretation and limiting the direct extrapolation of results across experimental models.

While the potential expression of nicotinic acetylcholine receptors in cortical BCs remains unresolved, with conflicting evidence in the literature (*53–55*), our findings demonstrate that ChCs undergo significant cholinergic modulation. One of the most striking observations is that direct activation of nAChRs, either pharmacologically (via carbachol or DMPP) or optogenetically (via BF cholinergic axon stimulation), evokes an inward current in ChCs (figure S2 and figure 3E-F). These responses persist under synaptic blockers and are abolished by MMA, a selective nAChR antagonist, demonstrating a direct action of ACh on ChCs. The primary nicotinic receptors in the cortex are homopentameric α7 and heteromeric α4β2 receptors. Our data show that α7-containing nAChRs play a minimal role in the cholinergic modulation of ChCs, as only a small fraction of ChCs express the α7 subunit (figure 2D), and MLA blockade does not significantly affect the inward current observed in electrophysiological recordings (figure 2G-H). In contrast, we detect robust β2 subunit expression in most ChCs (figure 2C), with a marked reduction in inward currents following blockade with DhβE (figure 2F). Interestingly, our results suggest the presence of α3-containing heteromeric nAChRs, likely α3β4, in cortical ChCs. While this receptor has been well-characterized in other brain regions(*56*), it has been less frequently reported in the cortex, although recent transcriptomic studies provide evidence for the presence of α3 in the mouse cortex (*38*, *46*). Despite the small fraction of cells expressing α7 or α3, the prominent α3 signal in FISH experiments (figure 2E) and the significant reduction in inward currents with tropinone (figure 2I) imply that heteromeric α3β4 receptors may play a more prominent role in the cholinergic modulation of ChCs than α7. Notably, α4-containing nicotinic receptors, rather than α7, are essential for the normal development of ChC axonal arborization (*37*).

In the cortex, acetylcholine operates through both synaptic and volume transmission via nicotinic receptor subtypes with complementary kinetic properties. α7 nAChRs respond rapidly to transient, high acetylcholine concentrations at anatomically defined synapses, generating fast inward currents that provide precise feedforward control of network activity. In contrast, α4β2 nAChRs act over a broader spatial and temporal scale: at synaptic and perisynaptic sites they mediate slower postsynaptic responses to sustained ACh availability, while in the extracellular space they participate in volume transmission, enabling tonic gain modulation of neuronal excitability (*56–58*).

Within this framework, the mode of cholinergic transmission—synaptic versus volume-mediated—critically determines how ChCs influence cortical computation. Volume transmission, with its slow, spatially diffuse action, is well aligned with the properties of high-affinity α4β2-containing nAChRs expressed by ChCs and can define a global set-point for cortical excitability, adjusting network tone and signal-to-noise ratio over extended timescales. Under sustained cholinergic levels—such as those prevailing during wakefulness or heightened arousal—ChCs would integrate this tonic signal and gradually modulate their own excitability, consistent with the predominance of β2- and α4-containing nAChRs in these cells.

By targeting the AIS, ChCs can dampen or synchronize PN firing depending on the timing and context of inhibition (*4–6*). Our findings refine this view by showing that elevated cholinergic tone, particularly during high-arousal states, robustly excites ChCs via nAChRs (figure 6). Electrophysiological recordings during physostigmine application revealed a robust depolarization of the resting membrane potential accompanied by a decrease in rheobase (figure 4A-B), thereby enhancing ChC responsiveness to near-threshold inputs. This effect was strongest at low current intensities and progressively diminished as input strength increased (figure 4C), indicating that cholinergic drive primarily lowers the recruitment threshold rather than simply scaling maximal firing rates. Such a mechanism would bias local inhibition toward more temporally precise control at the AIS, effectively gating PN output during behavioral states that demand rapid sensorimotor integration or sustained attentional engagement (*32–34*, *49*). Notably, this effect was abolished by nicotinic receptor blockade (figure 4D-F), confirming that the physostigmine-induced increase in cholinergic tone enhances ChC excitability via nicotinic mechanisms.

ChCs exhibit heightened activity during locomotion across several brain regions, including the hippocampus, visual cortex, and M2 (*17*, *18*, *20*, *36*). In the visual cortex, ChC activity closely tracks pupil dilation during locomotion, supporting their preferential recruitment during high-arousal states (*17*, *20*). Extending these findings, our data show that M2 ChCs correlate strongly with both locomotion and pupil fluctuations (figure 5D-H), indicating sensitivity to global arousal dynamics rather than purely motor output. These observations align with the established role of ACh in shifting cortical state from quiet wakefulness to active engagement, with pupil diameter serving as a reliable proxy for neuromodulatory drive (*35*, *47*, *49*).

The BF—the primary source of cortical acetylcholine—dynamically adjusts network excitability as animals transition between behavioral states (*25*, *28*, *29*). Our optogenetic experiments confirm that this system functionally innervates M2 ChCs (figure 3), supporting a model where tonic cholinergic signaling defines a state-dependent excitability window. Thus, these interneurons emerge as a dynamic interface where global neuromodulation and AIS-targeted inhibition converge: while cholinergic tone dictates the timing of their recruitment, their activation implements a precise gating of cortical output (Jung et al., 2023, 2022; McGinley et al., 2015a, 2015b; Vinck et al., 2015; Yang and Kwan, 2021). This framework aligns with recent evidence from the medial prefrontal cortex, where ChCs encode stimulus salience to drive associative learning by integrating long-range inputs from the anterior insula and paraventricular thalamus (*60*). In this context, tonic nicotinic signaling functions as a state-dependent gain control that sets the activation threshold for ChCs. By modulating ChC excitability, this cholinergic drive enables the broadcasting of salience information to pyramidal ensembles, effectively linking global arousal with learning and attention.

While our study offers robust evidence for cholinergic regulation of ChCs, several critical questions remain unresolved. First, although we identified key subunits of nicotinic acetylcholine receptors, the precise role of α3 subunits and other less common subunits demands further investigation to clarify receptor composition and kinetics fully. Second, although in vivo nicotinic blockade establishes a causal link between nicotinic signaling and changes in ChC engagement, this approach impacts not only ChCs but also other cell types expressing nAChRs, which may, in turn, influence ChCs through synaptic interactions. Therefore, more precise manipulations targeting nicotinic inputs specifically to ChCs are needed. Additionally, the potential integration of noradrenergic or serotonergic inputs remains to be explored. Finally, while we have examined the impact of cholinergic blockade on ChC activity, it will be crucial to assess how alterations in nicotinic modulation of ChCs could disrupt the proper functioning of the cortical network.

In summary, our results reveal that ChCs in M2 are subject to pronounced and direct cholinergic modulation via β2- and α3-containing nAChRs, in a manner consistent with engagement by nicotinic volume transmission. These findings support a functional model in which tonic cholinergic tone sets the excitability window of ChCs, determining when they are recruited to exert AIS-targeted, projection-specific inhibition over PN ensembles during high-arousal states. Future studies dissecting how distinct neuromodulatory pathways and receptor subtypes converge onto ChCs—and how perturbations of nicotinic signaling alter downstream PN activity—will be crucial to test these predictions and to clarify the contribution of ChCs to circuit-level computations in both physiological and pathological conditions.

## Methods

All procedures related with animal investigation were performed in accordance with the European Union Directive 2010/63/EU and Spanish law (R.D. 53/2013 BOE 34/11370-420, 2013) for the use and care of laboratory animals and approved by the competent Spanish entity (“Consejería de Agricultura, Pesca y Desarrollo Rural. Dirección General de la Producción Agrícola y Ganadera”) with authorized project number 16/12/2020/146. All mice were maintained in the university animal facility (Centro de Experimentación Animal Óscar Pintado) of the Centro de Investigación, Tecnología e Innovación de la Universidad de Sevilla (CITIUS) on a 12h-light/12h-dark cycle and housed in standard cages with unrestricted access to food and water.

### Animals

Both male and female mice (∼2 months old) were used for all experiments. The following transgenic lines were obtained from The Jackson Laboratory: *Vipr2*^Cre^ (strain 031332), *Pvalb*^Flp^ (strain 022730), *Ai65D* (strain 021875) and *Chat*^Cre^ (strain 006410), *PV*^Cre^ (strain 008069) and *Ai14* (strain 007914). To genetically label ChCs with a tdTomato reporter, *Vipr2*^Cre^:*Pvalb*^Flp^ mice were crossed with *Ai65D*, following the breeding strategy described by Tasic et al., 2018, generating the *VipR2-Pvalb-Ai65D* mouse line. To compare ChCs with parvalbumin-expressing cells, *PV*^Cre^ and *Ai14* mice were crossed to generate the *PVCre*-*Ai14* mouse line. Finally, in optogenetic experiments aimed at targeting cholinergic cells with ChR2, a quadruple transgenic line (*VipR2-Pvalb-Ai65D-ChAT*) was generated by crossing *Vipr2*^Cre^:*Pvalb*^Flp^ mice and *Ai65D*-*Chat*^Cre^ mice. Genotyping was performed using PCR and according to supplier protocols. All experimental mice were heterozygous for their respective transgenes.

### Tissue fixation and slices preparation

Mice were deeply anesthetized with 2% tribromoethanol (0.15 ml / 10 g) and then transcardially perfused with 0.9% saline solution, followed by 4% paraformaldehyde (PFA) in 0.1 M phosphate buffer (PB, pH 7.4). The brains were removed, post-fixed in 4% PFA overnight at 4°C, and then cryoprotected in 30% sucrose in PB containing 0.01% sodium azide until sectioning. Brains were subsequently frozen with O.C.T. compound (Tissue Tek) and then serially sectioned on a cryostat (Leica CM1950) into 40 µm-thick coronal slices. These slices were stored at -20°C in 50% glycerol in phosphate buffered saline (PBS, pH 7.4). In some experiments (figures 3 & 5), following electrophysiology, 300 µm-thick slices were fixed in 4% PFA overnight at 4°C, then cryoprotected and resectioned into 40 µm-thick coronal slices.

### Immunofluorescence

Free-floating brain slices were washed with PBS and then blocked with 3% normal serum and 0.3% Triton X-100 in PBS for 45-60 min. Next, slices were incubated overnight at 4°C with primary antibodies in the same blocking solution. After washing with PBS, slices were incubated for 2 hours at room temperature with appropriate secondary antibodies (Goat or Donkey Alexa Fluor 488, 647 or Rhodamine Red-X; 1:500, Jackson Immunoresearch) diluted in 1.5% normal donkey serum and 0.3% Triton X-100 in PBS. Finally, slices were washed and mounted on slides using FluorSafe Reagent (Cat#: 345789, Merk Millipore). The primary antibodies used were chicken CPCA-mCherry (1:1000, EnCor Bio, Cat#: CPCA-mCherry), rabbit Anti-parvalbumin (1:1000, Swant, Cat#: PV27), mouse Anti-parvalbumin (1:1000, Swant, Cat#: PV235), rabbit Anti-Phospho-IκBα (1:1000, Cell Signaling Technology, Cat#: 2859), goat Anti-ChAT (1:500, MerckMillipore, Cat#: AB114P), rabbit Anti-GFP (1:500, Synaptic Systems, Cat#: 132002) and chicken Anti-GFP (1:1000, Aves Labs, Cat#: #GFP-1010). An antigen-retrieval protocol was performed for experiments involving the goat anti-ChAT antibody before immunofluorescence. Briefly, slices were washed with PBS and incubated in sodium citrate solution (pH 6.0; Sigma) at 80°C in a water bath for 30 min. After rinsing with PBS, slices were blocked with 2% nonfat milk and 0.3% Triton X-100 in PBS for 30 min. Following another rinse, slices were blocked again in 3% normal serum and 0.3% Triton X-100 in PBS for 45–60 min, then incubated for 48 hr at 4°C with goat anti-ChAT. Finally, slices were incubated for 2 h at room temperature with the corresponding secondary antibodies and mounted on slides using FluorSafe Reagent.

### Fluorescent *in situ* RNA hybridization

Brains were obtained as described in *Tissue fixation and slices preparation*. They were then frozen with O.C.T. compound and serially sectioned on a cryostat (Leica CM1950) into 10 µm-thick coronal slices. Sections were mounted on SuperFrost Plus adhesion slides (J1800AMNZ, Epredia) and stored at -80°C until use. The *in situ* hybridization was performed using the ViewRNA kit (ViewRNA technology, Invitrogen, ThermoFisher Scientific, QVT4600C) for fixed frozen tissue, following the manufacturer’s instructions. All incubations were performed using an ACD HybEZ II Hybridation System. To minimize tissue damage, slides were gradually warmed from -20°C to 4°C and then to room temperature. A hydrophobic barrier was drawn around the sections with a hydrophobic pen (ImmEdge). Target probes used were Chrna3 (ID #VF6-10407-VC), Chrna7 (ID #VB6-3198060-VT) and Chrnb2 (ID #VB6-19634-VT). Negative (VF6-10407-VC) and positive (Actb, VB6-12823-VT) control probes provided in the kit were also used. A type 6 label probe (Alexa Fluor 647, Cy5 fluorochrome) was used for detection, and DAPI was added in the final steps to stain nuclei. To enhance the identification of ChCs, an immunofluorescence protocol against tdTomato using mCherry-CPCA was performed as previously described.

### AAV design and production

For cloning of AAV-hSynP-Flex-Tdtomato, each component expression cassette (human Synapsin Promoter–LoxP–tdTomato–LoxP) was amplified individually using custom-designed primers. The fragments were then assembled via overlapping PCR using Q5 High-Fidelity DNA Polymerase (M0491S, New England Biolabs). The resulting product was cloned into a pre-existing adeno-associated virus (AAV) vector containing ITRs (#18916, Addgene) at the NdeI and HindIII restriction sites. Enzymes and reagents included NdeI (R0111S), HindIII (R3104S), DpnI (R0176L), Antarctic Phosphatase (M0289), and T4 DNA Ligase (M0202S) (all from New England Biolabs).

AAV vector production was performed by the Viral Vector Unit at “Centro Nacional de Investigaciones Cardiovasculares” (CNIC). For encapsulation of tdTomato, the plasmid pUCmini-iCAP-PHP.eB (plasmid #103005, Addgene) was employed to provide the PHP.eB capsid. Viral titers were quantified by qPCR using primers targeting the woodchuck hepatitis virus post-transcriptional regulatory element (WPRE).

### Acquisition and image analysis

Confocal imaging was performed on a Nikon A1R+ microscope with a 40x air objective lens or 20x and 60x glycerol immersion objectives (Nikon), depending on the experiment. For RNA hybridization studies, confocal images were acquired using a Leica Stellaris 9 microscope, employing a 20x glycerol immersion objective lens (Leica) and 3x digital magnification. All images presented in the figures were acquired via confocal microscopy, except for those in figures 5 and 6, which were obtained using a two-photon microscope. Image analysis was conducted in Fiji software (*61*), with the experimenter blinded to the genotype to ensure unbiased data processing.

#### IMARIS analysis

The images were analyzed using IMARIS 9.6 software (Oxford Instruments) to reconstruct Chandelier cells (ChCs) based on tdTomato labeling and the axon initial segments of pyramidal cells using p-IkBa immunofluorescence. Confocal images were acquired using a 20x objective to capture complete images of individual, isolated ChCs, facilitated by their relatively low abundance. The Filament Tracer tool was employed to reconstruct the axonal and dendritic branches of tdTomato-positive ChCs. To perform three-dimensional reconstructions of ChC cartridges and pyramidal AIS, confocal images were obtained using a 60x objective, and the Surfaces-Surfaces contact tool was applied (available in: https://imaris.oxinst.com/open/view/surface-surface-contact-area). Briefly, the analysis quantifies contact surfaces within defined regions of interest (ROIs), ensuring consistency in volume between images. Manual selection of proximal and distal ROIs targeted AIS rods and ChCs cartridges. Segmentation parameters, such as a maximum diameter of 0.6 µm for the green channel, and manual thresholding ensured accurate representation of structures. Using the Surface-Surface Contact tool, AIS rods (primary surface) and cartridges (secondary surface) were analyzed to determine their overlap, visualized as a voxel-thick unsmoothed surface above the primary structure.

### Stereotactic viral infection and window implantation

#### Stereotactic AAV injections

Two-month-old mice were anesthetized with isoflurane (5% for induction, 1.5-2% for maintenance in oxygen) delivered at a constant flow rate. The scalp was shaved, and carprofen (0.5 mg/mL), a nonsteroidal anti-inflammatory drug, was administered subcutaneously prior to surgery. Mice were then placed on a motorized stereotaxic frame (World Precision Instruments), and ophthalmic ointment was applied to prevent corneal drying. The body temperature was maintained at 37°C using a thermostatically controlled heating pad (Physitemp TCAT-2LV). The scalp was disinfected with betadine and 70% ethanol, and an incision was made to expose the skull. After cleaning the skull with cortex buffer (NaCl 125 mM, Glucose 10 mM, KCl 5 mM, HEPES 10 mM, MgSO4 2 mM, CaCl2 2mM), the stereotactic coordinates were marked.

For viral injections targeting L2/3 of the M2 cortex, a small burr hole was drilled at the following coordinates (relative to bregma): anteroposterior (A/P) +2.8 mm, mediolateral (M/L) +1 mm, dorsoventral (D/V) −0.25 mm from the brain surface. For BF injections, three target areas were used: the nucleus of the vertical limb of the diagonal band (VDB; A/P +0.8 mm, M/L +0.5 mm, D/V −4.5 mm), the nucleus of the horizontal limb of the diagonal band (HDB; A/P 0 mm, M/L +1.5 mm, D/V −5.1 mm), and the nucleus basalis of Meynert/substantia innominata (NBM/SI; A/P −0.35 mm, M/L +1.6 mm, D/V −4.55 mm). The following viruses were injected as indicated in the figures: hSyn-tdTomato-FLEX or AAV1-EF1a-double-floxed-hChR2(H134R)-EYFP-WPRE (Addgene, Catalog #20298). The virus was loaded into a pulled glass micropipette, backfilled with mineral oil. The flow rate (5 nl/s) was controlled by a nanoinjector (Nanoject III, Drummond), delivering a total volume of 150-200 nl for M2 injections and 450 nl for each BF injection site. The pipette remained in place for 10 minutes before and after the injection to prevent backflow. The burr hole was sealed with silicon sealant (Kwik-cast, World Precision Instruments), and the incision was closed using skin adhesive (3M Vetbond). Mice were monitored until they recovered from anesthesia. For 3 days following surgery, mice received daily injections of carprofen (0.5 mg/ml).

#### Stereotactic AAV injections with cranial window

For the cranial window surgery, the scalp was removed after disinfection and cleaned with cortex buffer solution. The connective tissue was carefully excised with a scalpel to fully expose the skull, followed by the application of a thin layer of cyanoacrylate. The posterior portion of the skull was covered with black dental adhesive (Orthocryl, Dentaurum), and a custom-made headbar was affixed on top. A 2 mm diameter craniotomy was created around the M2 stereotactic coordinates to expose the underlying brain. The viral construct AAV1-Syn-Flex-GCaMP6s-WPRE-SV40 (Addgene, Catalog: #100845-AAV1) was injected as previously described.

To prepare the cranial window, a 3 mm diameter round glass cover (Warner Instruments, Cat#: 64-0720) was joined with a 1 mm diameter square glass cover (handmade with a diamond-tipped blade) using UV-curable optical adhesive (Thorlabs, NOA61). This assembled window was then implanted into the craniotomy and sealed with cyanoacrylate. The remaining skull surface was covered with dental adhesive. Mice were monitored until they recovered from anesthesia. For 3 days following the surgery, mice received daily injections of carprofen (0.5 mg/ml).

#### In vivo cholinergic blockade

Nicotinic antagonists (MMA and MLA) were dissolved in warm cortex buffer, filtered, and then injected into the cortex using a glass micropipette and a nanoinjector (Nanoject III, Drummond). During the initial surgery and AAV injection, a small hole was drilled through each coverslip intended for the cranial window. This hole was sealed with Kwik-Sil and covered with acrylic to prevent desiccation and infection.

Imaging sessions occurred three weeks after the AAV injection. The Kwik-Sil plug was removed, and GCaMP6-expressing ChCs were imaged for 8–10 minutes. Following this, the mouse was placed under a dissection microscope, where the glass micropipette containing the filtered nicotinic antagonists was inserted into the cortex via the pre-drilled hole, preserving the integrity of the cranial window. After drug delivery, the mouse was repositioned under the two-photon microscope, the same neurons were located, and their activities were imaged again for another 8–10 minutes to assess the effects of the antagonists(*62*)

### In vitro electrophysiology

#### Preparation of brain slices

Mice were anesthetized with 2% tribromoethanol (0.15 ml/10 mg) and decapitated. Brains were dissected-out and submerged in ice-cold cutting solution containing (in mM) 222 sucrose, 11 D-glucose, 3 KCl, 1 NaH_2_PO_4_, 26 mM NaHCO_3_, 0.5 mM CaCl_2_ and 7mM MgCl_2_ (osmolality 305-315 mosmol/kg), pH 7.4, when bubbled with carbogen (5% CO2 - 95% O2). 300 μm thick coronal slices were cut using a Leica VT 1200S vibratome (Leica Biosystems, Wetzlar Germany). Slices were allowed to recover for 25 min at 35°C in artificial cerebrospinal fluid (ACSF) containing (in mM) 124 NaCl, 10 D-glucose, 2.5 KCl, 1.25 NaH_2_PO_4_, 26 mM NaHCO_3_, 2.5 mM CaCl_2_, 2mM MgCl_2_ and 4 sucrose (osmolality 305-315 mosmol/kg), pH 7.4, when bubbled with carbogen (5% CO2 - 95% O2). They were subsequently kept at room temperature for 30-60 min before recording.

#### Electrophysiological recordings, data acquisition and analysis

Slices were transferred to a recording chamber installed in an upright microscope (Nikon Eclipse Ni-E). Recordings were carried out at 33 ±1C° with continuous perfusion at a rate of 2-3 ml/min. Cells were visualized with a 40X water immersion objective lens (Nikon, NA 0.8, IR 40×/0.80 W) with infrared light source (Thorlabs, M940L3) and Dodt-Gradient-Contrast System (Luigs & Neumann) and a USB 3.0 monochrome camera (TheImagingSource, DMK 23U445). ChCs were identified as tdTomato-positive neurons using a high-power multi-LED (pE-300white, CoolLED). Whole-cell patch-clamp recordings were carried out using an EPC10 amplifier (HEKA). Patch pipettes were pulled from borosilicate glass (Science Products GmbH, Cat#: GB150F-8P) using a DMZ Universal Puller (Zeitz Instruments). After filling the pipettes with intracellular solution containing (in mM) 120 K-gluconate, 10 KCl, 10 phosphocreatine, 10 HEPES, 0.1 EGTA, 2 Mg-ATP, 0.3 Na_2_-GTP (pH 7.25, 280-290 mosmol/kg) for all recordings, their resistances were 2.5-3.5 MΩ. Giga seals (>1GΩ) were always obtained before break-in. Throughout voltage-clamp recordings, whole-cell capacitance and series resistance were monitored, and series resistance was compensated by 70% (lag 10 µs). In all experiments, the neurons were discarded whether resistance increased more than 50% or exceeded 20 MΩ. For local application of 1,1-Dimethyl-4-phenyl piperazine iodide (DMPP), a glass pipette delivered a 20-ms, 4-psi puff to the dendritic region near the soma of ChCs, controlled by an Openspritzer (*63*). Bridge-balance and capacitances were fully compensated in current clamp recording.

Recordings were acquired using PatchMaster software (HEKA) with a low-pass filter at 2.9 kHz and digitized at 20 kHz. A D400 Multi-channel 50Hz/60Hz mains noise eliminator (Digitimer) was used. In current-clamp recordings, input resistance (Rin) was determined by injecting current pulses (10 pA increment, from –50 pA to +50 pA, 500 ms duration, 1 Hz) estimating the slope of the voltage-current plot. The membrane time constant (τ) was obtained by recording the membrane response to - 50 pA current pulses (500 ms duration, 1 Hz) and fitting the averaged record of 10 sweeps to a single exponential curve. The rheobase was the minimum injected current (500 ms pulse duration at 1 Hz) to generate an action potential with 50% likelihood. Firing properties were obtained during current injection steps (20 pA increments, 500 m) and represented as frequency-intensity (FI) plots. Action potential (AP) parameters were calculated from the rheobase recording. The AP threshold was calculated as the value of the membrane potential at which the first derivative surpassed 20 V/s. Recordings were analyzed with the software Stimfit (*64*).

#### Optogenetic stimulation

To investigate BF cholinergic inputs to ChCs, we patched ChCs located in layer 2/3 of M2 that were surrounded by a high density of eYFP-positive cholinergic axons. Optogenetic stimulation of BF cholinergic axons was delivered with 5 s pulses of blue light (2.5 mW/mm², 450 nm) using a high-power multi-LED system (pE-300white, CoolLED). Postsynaptic currents were recorded in voltage-clamp mode at a holding potential of −70 mV. To record monosynaptic transmission, we applied to the bath TTX (500 nM) to block action potential–dependent synaptic release and 4-aminopyridine (4-AP, 500 µM) to facilitate transmitter release from BF axons and thereby enhance the detectable response at monosynaptic connections (*65*).

### Calcium signaling via two-photon imaging

#### Data collection and processing

Mice were head-fixed on a floating spherical treadmill, trained to balance and run during daily 20-minute sessions for five consecutive days before experimental procedures began. Imaging sessions were initiated 18–21 days post-AAV injection, upon confirmation of GCaMP6 expression in the M2 region. A resonant scanning two-photon microscope (Neurolabware, Los Angeles, CA) equipped with a pulsed Ti:sapphire laser (Chameleon Discovery, Coherent) and controlled by Scanbox acquisition software (Scanbox, Los Angeles, CA) was used for imaging. A 16x water immersion objective (Nikon, 0.8 NA, 3 mm working distance) was employed at a zoom level of 1.4–1.7x. Laser excitation wavelengths were set to 920 nm for GCaMP6 signals and 1100 nm for tdTomato. The microscope operated at a frame rate of 15.6 Hz (512 lines using a resonant mirror at 8 kHz).

Imaging planes were chosen based on cell density and the extent of viral infection, targeting depths of 150–300 μm. GCaMP6 fluorescence signals were recorded for 8–10 minutes per plane, while tdTomato signals were captured over 30 seconds to facilitate post hoc identification of Chandelier cells.

Experiments were conducted in the dark, illuminated with infrared light for visibility. Pupil size and ball movement were recorded concurrently using two cameras, each fitted with a 740 nm long-pass filter: a Dalsa M1280 at 30 fps for pupil monitoring and a Dalsa M640 at 60 fps for tracking ball movement. Both locomotion and eye movement data were synchronized with microscope frame acquisition. To pre-constrict the pupil, during the imaging session a small, non-dispersing light source was aimed at the contralateral eye.

Calcium imaging data were processed using MATLAB scripts provided by Scanbox, which included motion correction, segmentation, and signal extraction. Comparable results were achieved using the Suite2P pipeline (*66*). MATLAB scripts from Scanbox were also used to analyze pupil signals and ball movement.

#### Data analysis

An analysis of calcium signals from ChCs, pupil dilation, and ball movement was performed using MATLAB. For the data in figure 5, correlation coefficients were computed to assess the relationship between each ChC’s activity and both locomotion and pupil diameter. GCaMP6s signals were classified as either tdTomato-positive or tdTomato-negative by aligning the tdTomato and GCaMP6 masks. For the data in figure 6, a cross-correlation analysis was carried out to evaluate the relationship between motion and ChC activity. Additionally, changes in normalized amplitude were measured before and after the administration of nicotinic antagonists.

### Statistical analysis

All statistical analysis was made using Past software (*67*). The assessment of statistical significance was guided by tests for homogeneity of variance (Levene’s test) and normality (Shapiro-Wilk test) applied to the compared samples. Depending on the distribution characteristics, statistical significance was determined using either Student’s t-test or Mann-Whitney U test for pairwise comparisons. In situations where the datasets were paired, statistical significance was determined using paired Student’s t-test or Wilcoxon signed-rank test. For repeated measures, we employed ANOVA followed by Tukey’s post-hoc pairwise comparisons, or the Friedman test followed by Wilcoxon signed-rank tests with Bonferroni correction to identify specific differences between conditions. Box plots illustrate the 25th to 75th percentiles, represented by the box, with the median shown as a horizontal line within the box. Whiskers represent the minimum and maximum values.

### Drugs and Solutions

All drugs and chemicals were from ThermoFisher Scientific (Waltham, MA), except for TTX (Latoxan Laboratory, France), CNQX, DL-AP5 and Picrotoxin (HelloBio, Princeton, NJ), Physostigmine, MMA, DHβE and MLA (TOCRIS, UK) and Atropine and Tropinone (Sigma-Aldritch, St. Louis, MI).

## Acknowledgements

We gratefully acknowledge the support and advice provided by R. Fernández-Chacón. We thank J. Lopez Barneo for carefully reading the manuscript and for providing insightful comments and constructive criticism that helped improve the paper. We also acknowledge the help with Imaris software by Nozha Borjini, the excellent technical assistance of C. Cabrera-Romero and G. Asensio Gómez, and the technical support of the staff at IBiS and the Centro de Investigación, Tecnología e Innovación de la Universidad de Sevilla (CITIUS). This project was supported by: grant PGC2018-095656-B-I00 funded by MICIU/AEI/10.13039/501100011033 (PGJC and JLNG) and by ERDF “A way of making Europe”, grant PID2021-123840NB-I00 funded by MICIU/AEI/10.13039/501100011033 and by “ERDF/EU” (PGJC and JLNG), grant US-1264432 funded by US/JUNTA/FEDER, UE (PGJC and JLNG), grant RyC-2016-19906 (PGJC), grant 2019/FTATIANA/FCHACON funded by Fundación Tatiana Pérez de Guzmán El Bueno (PGJC), PRE2019-087729 (EMM) funded by MICIU/AEI/10.13039/501100011033 and by “ESF Investing in your future”, grant RyC-2016-19906 funded by VI PPIT-US (PGJC), grant PREDOC_00594 funded by Junta de Andalucía (SRL) and SLB was supported by project PID2019-105530GB-I00 funded by MICIU/AEI/10.13039/501100011033.

## Author contributions

Emilio Martinez-Marquez, data collection, data analysis, reviewing/editing; Santiago Reyes-Leon, data collection, data analysis, writing, reviewing/editing; Flores García-Sanjosé, data collection and data analysis, Santiago López-Begines, methodology; Pablo Garcia-Junco-Clemente and Jose L. Nieto-Gonzalez, conceptualization, experimental design, data collection, interpretation, writing, review/editing and funding.

## Competing interests

Authors declare that they have no competing interests.

## Figures and figure legends

**Supplementary figure 1.**
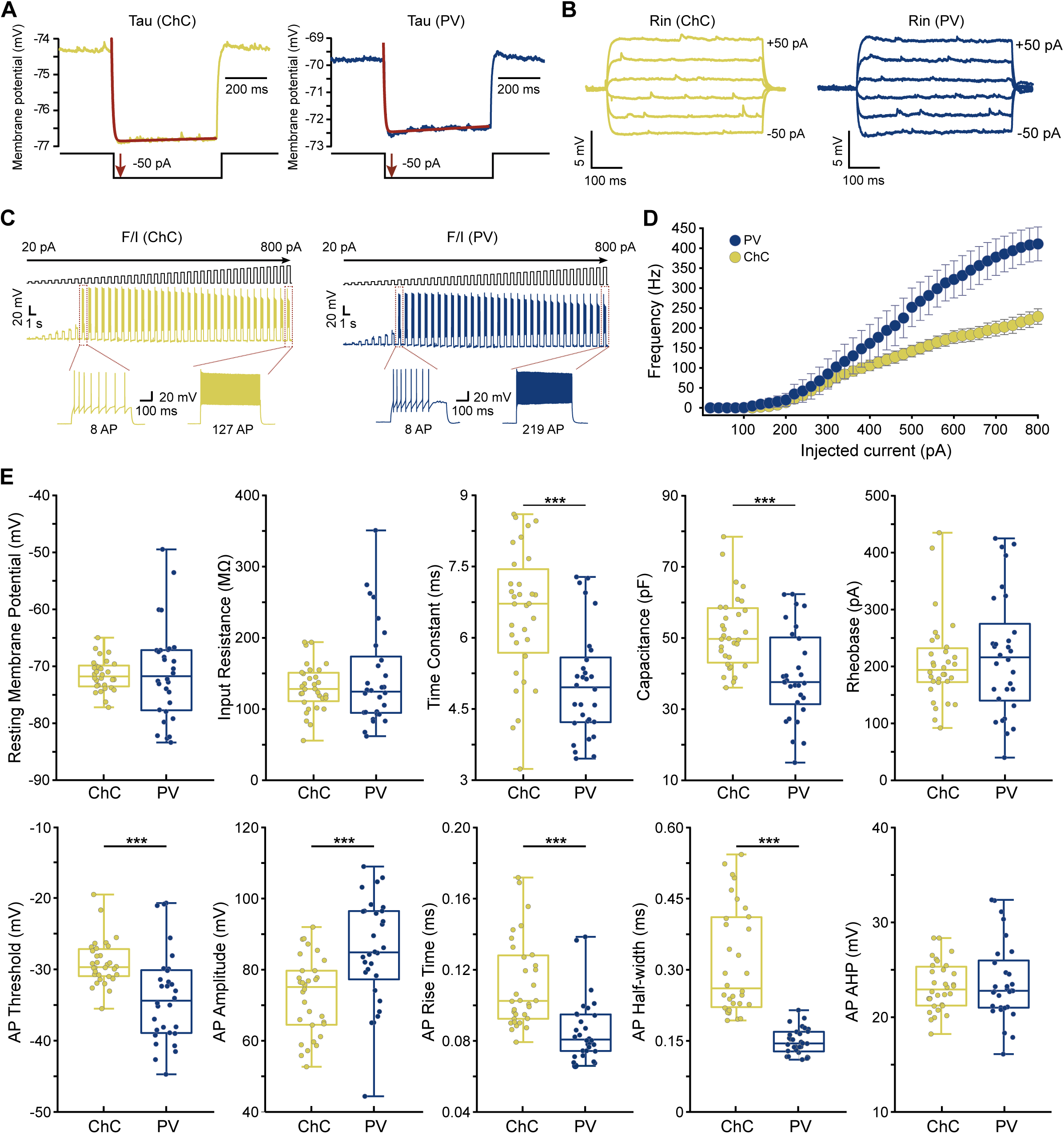
Comparative analysis of intrinsic membrane properties and firing dynamics in Chandelier versus Parvalbumin-expressing interneurons. **(A)** Representative traces of subthreshold membrane potential responses to a single −50 pA current injection step (top traces) and the corresponding injected current steps (bottom traces) for a ChC (left) and a PV cell (right). Exponential fitting of the voltage response (red) was used to determine the membrane time constant. **(B)** Representative traces showing subthreshold voltage responses to current injections ranging from −50 pA to +50 pA for a ChC (left, yellow) and a PV cell (right, blue). Input resistance was calculated from the steady-state voltage deflections. **(C)** Representative F/I traces from a ChC (left, yellow) and a PV cell (right, blue) illustrating how action potential firing frequency increases in response to increasing injected current steps. Current pulses ranging from 20 pA to 800 pA (top traces) are shown. Magnified examples of AP firing with low current and high current are provided to show characteristic spike phenotypes. **(D)** Plot of action potential frequency (Hz) as a function of injected current (pA) for ChCs (yellow) and PV cells (blue). Unpaired *t*-tests were performed at each current step, showing that PV cells exhibit significantly higher firing rates than ChCs at larger current intensities (n= 17-30 (4 mice) for ChCs and n= 21-30 (3 mice) for PV cells, depending on the depolarizing current step) (\**p* < 0.05, **p < 0.01, \*\*\**p* < 0.001 and *p< 0.0001). **(E) Summary of intrinsic and action potential properties.** Statistical comparison of electrophysiological parameters reveals distinct profiles for both cell types. ChCs exhibited a significantly longer membrane time constant and higher capacitance compared to PV cells, while no significant differences were observed in resting membrane potential, input resistance, or rheobase. Regarding action potential kinetics, ChCs showed a more depolarized AP threshold, along with a significantly lower AP amplitude and slower kinetics, characterized by longer AP rise times and wider AP half-widths than those of PV cells. AP afterhyperpolarization remained similar between groups. All data are presented as boxplots with individual data points overlaid (n= 32 ChCs, 4 mice); asterisks denote ***p < 0.001 by unpaired t-test.

**Supplementary figure 2.**
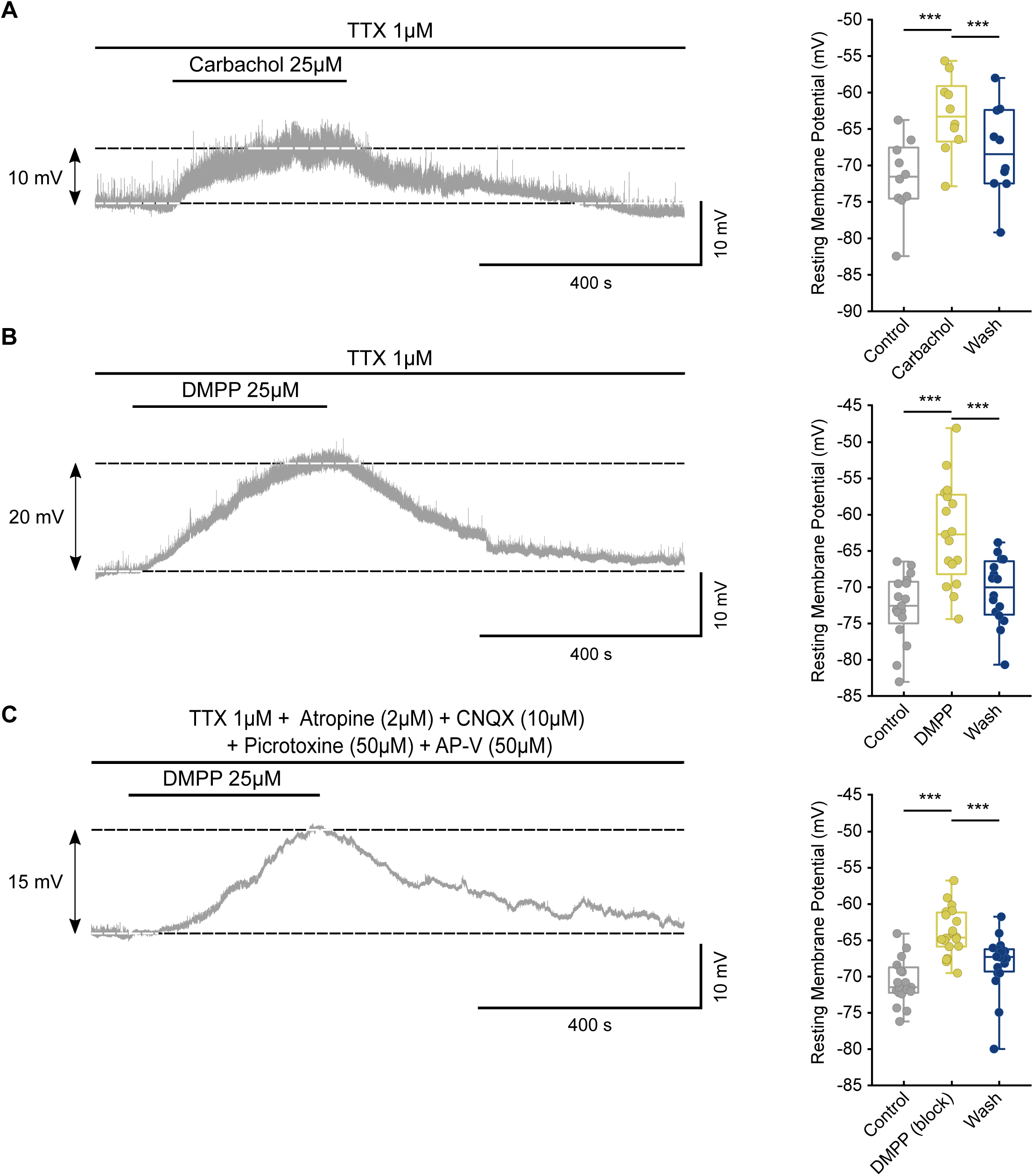
Cholinergic activation directly depolarizes Chandelier Cells in the secondary motor cortex. **(A)** Representative whole-cell patch-clamp recordings of ChCs in the M2 cortex showing a reversible depolarization of the resting membrane potential following bath application of the general cholinergic agonist carbachol (25 µM) in the presence of TTX (1 µM). The accompanying box plot quantifies the significant depolarization and subsequent washout (***p < 0.001, repeated measures ANOVA with Tukey’s post hoc test, n= 9 cells, 5 mice). **(B)** A similar depolarizing shift is elicited by the nicotinic agonist DMPP (25 µM), indicating the involvement of nicotinic acetylcholine receptors (nAChRs). Box plot summarizes the magnitude of the response and its reversibility (***p < 0.001, Friedman test with Wilcoxon post hoc and Bonferroni correction, n= 14 cells, 8 mice). **(C)** The DMPP-induced depolarization persists during the application of a pharmacological cocktail (TTX 1 µM, CNQX 10 µM, DL-APV 50 µM, picrotoxin 50 µM, and atropine 4 µM) to block voltage-gated sodium channels, ionotropic glutamatergic and GABAergic transmission, and muscarinic receptors. These results demonstrate that the nicotinic effect is independent of fast synaptic transmission and muscarinic signaling, suggesting a direct postsynaptic modulation of ChCs. (***p < 0.001, Friedman test with Wilcoxon post hoc and Bonferroni correction, n= 20 cells, 10 mice).

**Video 1 Simultaneous GCaMP6s imaging of Chandelier Cell somata and axonal boutons during spontaneous behavior.**

This video captures the calcium dynamics of ChCs in the M2 cortex recorded over approximately four minutes (original acquisition: 15.56 fps; playback accelerated to 120 fps). The high-resolution GCaMP6s signals allow for the visualization of activity not only in the cell bodies (somata) but also across their axonal cartridges and synaptic boutons. The neural activity is synchronized with real-time behavioral monitoring, including treadmill locomotion (ball movement) and pupil diameter. Notably, ChC calcium transients are highly correlated with arousal states, showing maximal intensity during active running bouts and periods of significant pupil dilation.

## Notes

### Competing Interest Statement

The authors have declared no competing interest.

### Summary of Updates

We have uploaded a revised version of our manuscript to incorporate new experiments and a refined conceptual framework. In this version, we introduce physostigmine experiments that demonstrate how acetylcholinesterase inhibition produces a gradual, sustained increase in chandelier cell excitability, fully dependent on nicotinic receptors, providing evidence for nicotinic volume transmission as a tonic regulator of these interneurons. We also clarify the contribution of specific nicotinic receptor subunits, strengthen the link between chandelier cell recruitment and behavioral markers of arousal, and streamline the narrative to emphasize their role as a bridge between global cholinergic arousal signals and local cortical output control. Finally, we have polished the text, updated the figures and legends, and expanded the discussion to better integrate our findings with current models of state‑dependent cortical computation.

